# Structural analyses of a heterodimeric SusCD complex captures intermediate states of maltooligosaccharide transport

**DOI:** 10.64898/2026.09.16.752182

**Authors:** Michael C. Cadigan, Nicole Rivera Fuentes, Wilhelm Salmen, Jacquelyn R. Roberts, Brandon Ruotolo, Melanie D. Ohi, Nicole M. Koropatkin

## Abstract

*Bacteroides* are abundant gut bacteria with diverse polysaccharide metabolizing capabilities encoded by co-transcribed polysaccharide utilization loci (PULs). The starch utilization system (Sus) is required for starch and α-glucan metabolism in *Bacteroides thetaiotaomicron* and has served as a model for PUL studies. However, the mechanism of maltooligosaccharide (MOS) transport and the architecture of the SusCD complex remain incompletely understood. Here, we used cryo-electron microscopy to capture multiple transport-relevant conformations of the SusD lipoprotein and the TonB-dependent transporter SusC in both unliganded and ligand-bound states. In the absence of ligand, SusCD adopts three conformations including an open state in which SusD is displaced from SusC, a state containing the SusC plug, and a state lacking density for the SusC plug. Three-dimensional variability analysis of the structure where SusD is displaced from SusC reveals continuous SusD mobility coupled to movement of the SusC N-terminus. The maltoheptaose-bound SusCD structure reveals two binding sites, one at the SusD-SusC interface and a second within SusC, indicating how MOS stabilizes a closed SusCD assembly. Notably, all structures show that SusCD is a heterodimer, unlike other Sus-like transporters that assemble into SusCD heterotetramers. Native mass spectrometry confirms this heterodimeric stoichiometry, and comparisons with other SusCD-like complexes suggest a structural basis for this distinct assembly of SusCD. Together, these data define substrate binding and intermediate transport states in a canonical PUL system and reveal architectural divergence among Sus-like carbohydrate transport complexes in *Bacteroides*.

## Introduction

The human gut microbiota, the community of microorganisms within the intestine, profoundly influences host health and disease (1–3). One key function of intestinal bacteria is the metabolism of dietary fiber, broadly defined as carbohydrates that are inaccessible to host enzymes (4). Anaerobic metabolism of these substrates produces short chain fatty acids, which shape host physiology by influencing processes such as the host immune response and energy harvesting (5). Members of the genus *Bacteroides* are dominant constituents of the gut microbiota and have an extensive capacity to degrade both host-derived carbohydrates, such as mucin, and dietary carbohydrates that pass through the intestine (6, 7). They organize this glycolytic potential within polysaccharide utilization loci (PUL), which are co-expressed gene clusters required to capture, breakdown, and transport specific polysaccharides or related classes of carbohydrates (7–9). Numerous studies have shown that the repertoire of PUL encoded by a given *Bacteroides* species define its metabolic niche in the gut and its ability to persist in the host despite dietary perturbations (8, 10, 11).

The starch utilization system (Sus) of *Bacteroides thetaiotaomicron* (*Bt*) was the first PUL to be characterized and is required for growth on starch and related α1,4, α1,6 glucans (12). The Sus encodes eight co-expressed genes (*susA-G and R*; **Fig 1**). At the cell surface, the lipoproteins SusD, SusE, and SusF bind starch, while the glycoside hydrolase family 13 (GH13) SusG cleaves it into maltooligosaccharides (MOS) (13–16). SusC is a TonB-dependent transporter (TBDT) that partners with SusD to transport MOS across the outer membrane into the periplasm (17).

**Figure 1.**
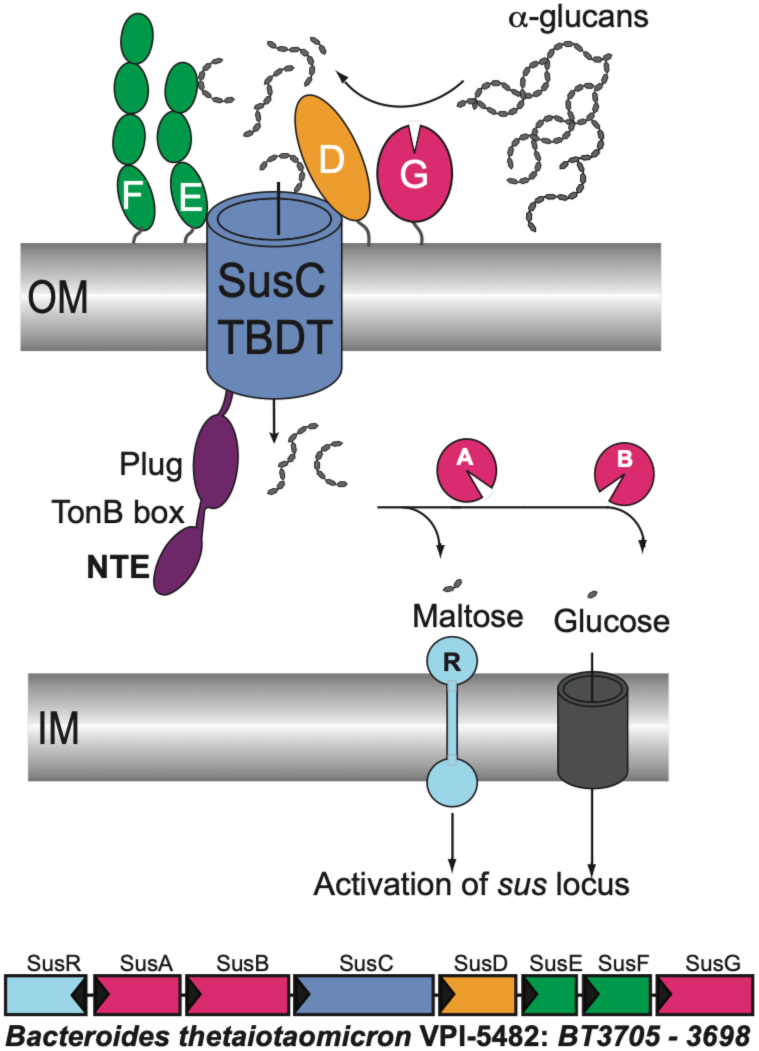
*Bacteroides thetaiotaomicron* (*Bt*) Starch utilization system (Sus). TonBdependent transporter (TBDT) SusC works with the outer membrane starch-binding lipoproteins SusD/E/F, and SusG, a glycoside hydrolase from family 13 (GH13). Periplasmic enzymes SusA (glycoside hydrolase family 13, GH13) and SusB (GH97) hydrolyze maltooligosaccharides to glucose before import into the cytoplasm via an unidentified transporter. SusR is a transcriptional regulator that spans the inner membrane and senses maltose in the periplasm to upregulate the operon. At bottom, organization of the Sus operon.

TBDTs are β-barrel proteins that contain a plug domain and a TonB box motif formed by residues in a β-strand (18). This motif mediates interactions with the inner membrane protein TonB which drives the import cycle (19). Like most glycan-targeting TBDTs in *Bacteroides*, SusC contains an N-terminal Ig-like domain, named the N-terminal extension (NTE), which is hypothesized to influence pairing to TonB (20). In the periplasm, the GH13 enzyme SusA and glycoside hydrolase family 97 (GH97) enzyme SusB further process MOS into glucose (21). SusR is an inner-membrane spanning transcriptional regulator that senses maltose, leading to the upregulation of *sus* expression that enables the capture of starch and starch-like structures (22) (**Fig 1**). More broadly, *Bacteroides* PUL encode similarly organized Sus-like systems that include a TBDT, lipoproteins, carbohydrate-degrading enzymes, and often a dedicated transcriptional regulator (9). A hallmark of these PUL is the presence of adjacent *susC*-like and *susD*-like genes (23).

Across PUL-encoded Sus-like systems, SusC-like and SusD-like proteins share sequence and structural homology, suggesting conserved function. Structures of SusCD-like complexes from members of the Bacteroidota, including *Bt*, *B. fragilis* and *Porphyromonas gingivalis*, show that the SusD-like lipoprotein opens and closes over the SusC-like TBDT(20, 24–27). In contrast to many vitamin or iron-targeting Proteobacterial TBDTs that function as monomers, the SusCD-like complexes form heterotetramers. The tetramer forms through dimerization of the SusC-like TBDTs, with a SusD-like protein anchored to each SusC-like protein (24–26). This arrangement appears to permit independent movements of each SusD-like “lid” over its partner transporter. Notably, the tetrameric assembly is stable and resistant to disruption by detergents used to extract these complexes from the outer membrane (20, 26). More recently, intact complexes of the SusCD-like tetramer together with additional cell-surface lipoproteins and/or enzymes from the Bt levan- and dextran-targeting PULs have been structurally characterized (27). The *Bt* genome has 88 PULs, and although genetic and transcriptomic studies have identified the target carbohydrate(s) for many of them, structural information is available for only a very small subset of the encoded proteins (28). As a result, it remains unclear whether Sus-like complexes assemble with the same stoichiometry or have distinct architectures.

Work on the *Bt* Sus, long considered the archetypal *Bacteroides* PUL, suggests that it may assemble differently from the *Bt* levan- and dextran-targeting systems described above. Co-immunoprecipitation with custom antibodies to the lipoproteins SusD or SusE failed to capture all cell surface Sus proteins, SusCDEFG, and sub-stoichiometric capture of SusC with SusD is consistent with partial dissociation between these components (29). Single molecule imaging further showed that the surface enzyme SusG, a lipoprotein, diffuses within the outer membrane, while lipoproteins SusE and SusF remain relatively fixed in position (29, 30). In addition, SusG enzymes from *Bt* and *B. ovatus*, despite having very different domain architectures, can be exchanged between these bacteria to restore function, suggesting that these surface enzymes do not need to work directly with the SusCD complex (31). Together, these observations support a model in which the *Bt* Sus assembles dynamically, in contrast to the more stable assemblies observed in other Sus-like systems systems (27). Many *Bacteroides* encode a dedicated Sus PUL, and α1,4, α1,6- linked glucans, such as pullulan and amylopectin, are the most widely consumed class of carbohydrates across *Bacteroides* type strains (32).

Here we present single particle cryo-electron microscopy (cryo-EM) structures of liganded and unliganded complexes of *Bt* SusCD. In all 3D reconstructions, *Bt* SusCD forms a heterodimer containing one copy each of SusC and SusD, rather than the heterotetramer observed in other SusCD-like systems. Native mass spectrometry analyses independently support the *Bt* SusCD stoichiometry observed by cryo-EM. Our maltoheptaose-bound *Bt* SusCD structure, SusCD_+M7_, captures two bound sugars within the SusC barrel and reveals ligand-associated conformational changes, including repositioning of SusD to close over SusC. Three-dimensional variability analysis (3DVA) of the unliganded SusCD complex further shows that when no substrate is bound, SusD appears to be minimally tethered to SusC, allowing it to adopt a dynamic wide-open conformation in relation to SusC. Further, in the unliganded SusCD complex the NTE of SusC is highly flexible, which may facilitate binding with the C-terminus of TonB during the import cycle. Together, our structures define protein-MOS interactions and transport-relevant conformational states, providing new mechanistic insights into starch specific import and expanding current models of carbohydrate uptake by *Bacteroides* Sus-like complexes.

## Results

### Cryo-EM reveals that unliganded SusCD is a conformationally flexible heterodimer

To determine the structure of SusCD, we solubilized the complex directly from *Bt* outer membranes using a two-step detergent extraction, adapted from the purification protocols for other *Bacteroides* SusCD-like complexes (20, 26). A C-terminal hexahistidine tag on SusD enabled affinity purification, followed by size exclusion chromatography (SEC) to obtain a biochemically homogeneous sample (**Fig S1A)**. Examination of the SusCD purification by SDS-PAGE did not show co-purification of additional Sus lipoproteins, such as SusE, SusF, and SusG (**Fig S1B**).

To structurally characterize the unliganded SusCD complex we vitrified the purified complex using ultrathin carbon grids and collected images on a 300 kV Titan Krios equipped with a Gatan K3 direct electron detector and a Bioquantum energy filter (**Fig S1C**). 2D classification of images of vitrified unliganded SusCD particles showed that the complex had clear secondary structural features and there appeared to be multiple views of the complex (**Fig S1D**). However, surprisingly, across all the 2D averages we could only detect what appeared to be a SusC monomer or a complex containing a single copy each of SusC and SusD. Even when the box size was increased to 480px (512.3 Å) during particle picking, which would generously accommodate a SusCD heterotetramer (SusC_2_D_2_) as reported for other *Bacteroides* SusCD-like transporters, we still only observed SusCD heterodimers (24–27).

Using single particle cryo-EM image processing methods, we determined three structures from this dataset of SusCD without ligand at global resolutions between 3.3-3.1 Å, which was sufficient to directly build atomic models for most of each protein directly into the maps (**Figs S2** and **S3, Table S1).** The three 3D structures include: 1) a map with density for SusCD and the SusC plug (SusCD_+Plug_) (**Fig 2A-D**); 2) a map of SusCD without density for the SusC plug (SusCD_NoPlug_) (**Fig 2E-H**); and 3) a map with density for only SusC (SusCD_Open_) (**Fig 2I-L**). As reported for previous SusCD-like structures (20, 24–26), density for the NTE, residues 26-104 preceding the TonB box, was not observed. The SusCD_+Plug_ and SusCD_NoPlug_ structures have a 1:1 SusC:SusD stoichiometry.

**Figure 2.**
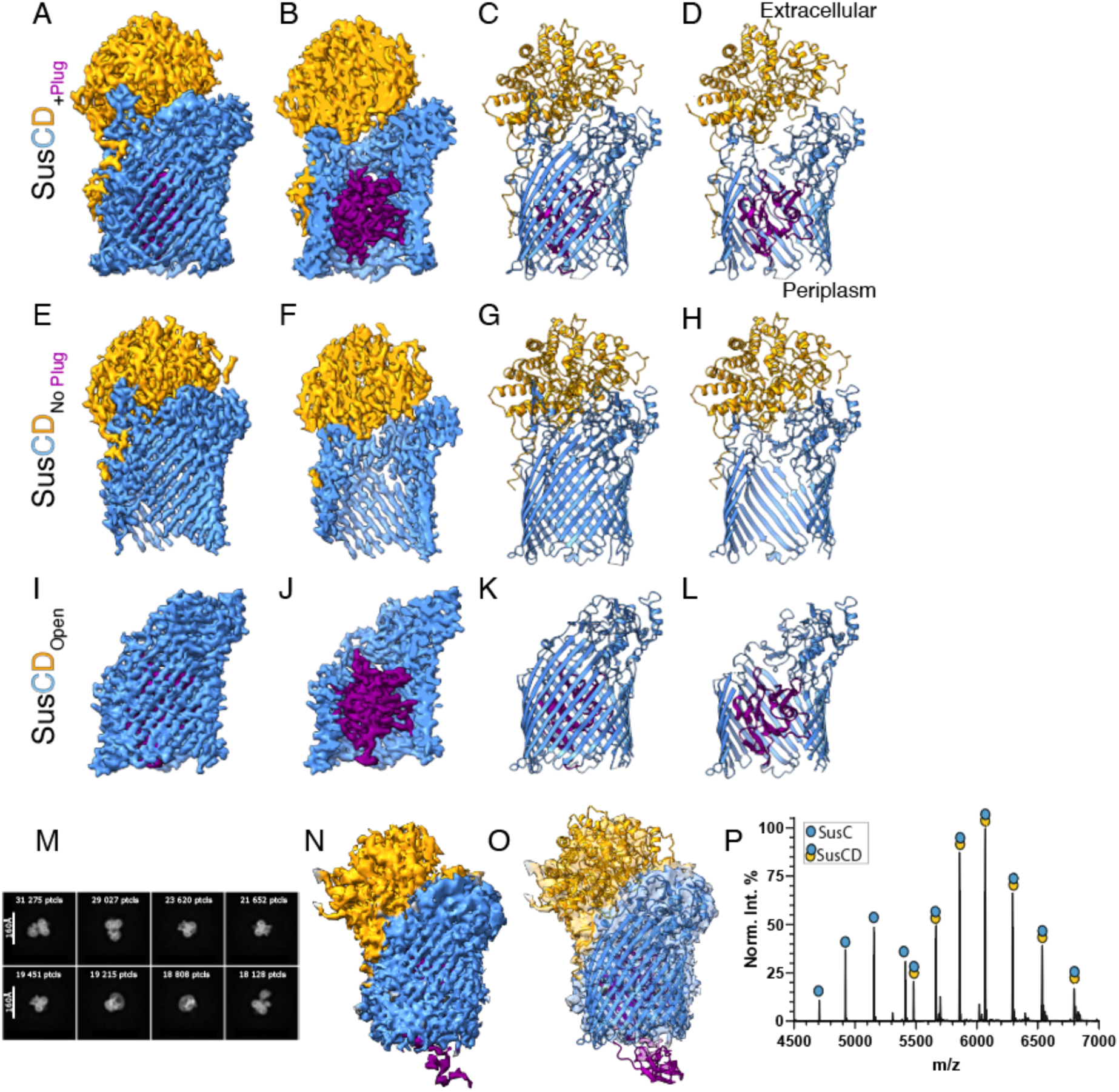
CryoEM structures and native MS demonstrate a SusCD heterodimer. A-D. 3.3Å resolution electron density map and protein model of SusCD^+Plug^ E-H. 3.1Å resolution electron density map and protein model of SusCD_NoPlug_, and I-L. 3.2Å resolution electron density map and protein model of SusCD_Open_. All maps and models shown in the same orientation with SusD in yellow, the SusC barrel in blue, and the SusC plug in purple. Panels B, D and equivalent panels of each series are clipped in the y-axis to show the interior of the SusC barrel. M. Representative 2D classes from SusCD_Open_ data processing. N. 4.4Å electron density map generated from a non-uniform refinement of the SusCD_Open_ volume from clustered 3D variability in CryoSPARC. O. Model of SusC, SusD, and predicted SusC_NTE_ domain rigid body fit into SusCD_Open_ 4.4A map. The SusC NTE is shown in purple. P. High-resolution native MS data of 3 μM SusCD demonstrating a heterodimeric complex.

### 3D variability analysis (3DVA) reveals flexibility of SusD and the SusC NTE in the open state

In the SusCD_Open_ structure, which lacked density for SusD, one possibility was that SusD had dissociated from SusC during vitrification. However, this structure also lacks density for two extracellular loops, EL7 and EL8, of SusC that may be a hinge upon which SusD pivots (see left side of SusC in **Fig 2A-H** vs **Fig 2I-L**), suggesting flexibility in this region of SusC and an alternative where SusD is present but flexibly tethered to SusC, causing its density to be averaged out during 3D reconstruction. Consistent with the latter possibility, several 2D class averages from this dataset appeared to show a diffuse, SusD-sized density extending from SusC at different angles (**Fig 2M**), suggesting that SusD flexibility relative to SusC accounts for the absence of well-resolved SusD density in the SusCD_Open_ structure.

To test this further, we performed 3DVA in cryoSPARC (**Fig S4**). This approach identifies structural modes corresponding to continuous conformational variability and can then partition particles into classes based on these modes (33). We analyzed SusCD particles suspected to contain SusD in the open conformation using three eigenvector modes (Modes 0, 1, and 2) each containing 20 total frames at a filter resolution of 5 Å. Frames in each movie represent shifts along the reaction coordinate from negative (frame 0), to positive (frame 19). Mode 0 (**Movie S1**) captured variability associated with SusD moving toward the closed position, where Mode 1 (**Movie S2**) captured variability associated with SusD moving toward the open position. Mode 2 primarily corresponded to variability in remnants of the unstructured detergent micelle that remained after particle subtraction and was therefore not considered further. Unexpectedly, as the reaction coordinate progressed from negative to positive, an additional density became visible on the periplasmic face of the SusC β-barrel, at the expected position of the SusC NTE (**Movie S1**).

We next used the cryoSPARC 3DVA clustering workflow to generate a map of the SusCD complex with SusD in the open conformation. Using the same 3DVA results shown in **Movie S1** and **S2**, particles were partitioned into 20 discrete clusters spanning the reaction coordinates, and clusters in which SusD adopted an open state were selected for additional local refinement using a whole complex SusCD mask. This approach yielded a ∼4.4 Å resolution map with density for SusD in a wide-open position, as well as density that we attribute to the SusC NTE (**Fig 2N,O, Fig S4**). The SusC and SusD models, together with an AlphaFold3 prediction of the NTE, fit well into this EM density map. The observation that NTE density was visible only after 3DVA supports the conclusion that this domain is conformationally flexible, consistent with prior studies indicating substantial NTE mobility (20, 26). Thus, 3DVA supports a model in which SusD remains flexibly tethered to SusC in the open state, while the SusC NTE samples multiple conformations on the periplasmic side of the β-barrel.

### Native mass spectrometry (MS) supports a 1:1 stoichiometry for the SusCD complex

A striking feature of the SusCD structure is the 1:1 stoichiometry of SusC and SusD. This contrasts with other SusCD-like complexes in *Bacteroides* which form core heterotetramers in which two SusC-like proteins dimerize and recruit two SusD-like partners (24–26). Our structural analysis showing that that SusC is a monomer in our structures makes it more similar to Proteobacterial TBDTs and the recently determined structures of *Bacteroides* vitamin B12 and iron-capturing TBDTs that form heterodimeric complexes with non-SusD nutrient-binding lipoproteins (34, 35).

To further assess SusCD stoichiometry, we used native MS, which preserves non-covalent interactions and enables direct measurement of protein complex oligomeric state. High-resolution native MS analysis showed that the predominant species in our sample was a SusC-SusD heterodimer with a deconvoluted mass of 169,897 ± 9 Da (**Fig 2P**), along with a minor population corresponding to SusC alone. The presence of free SusC was unexpected because the complex was purified using a C-terminal His_6_-tag on SusD, and because the SusD N-terminus (residues 25-43) makes extensive contacts with the exterior of the SusC barrel, burying ∼1200 Å^2^ of surface area. One explanation is that partial dissociation occurs after affinity purification, with some SusD lost during size exclusion chromatography.

### SusD engages the SusC β-barrel through a conserved hinge-like interface

Like other TBDTs, SusC folds into a 22-stranded β-barrel with 11 extracellular loops (ELs). SusD interacts with the SusC acting as a “lid” over the extracellular side of the SusC β- barrel. In both the SusCD_+Plug_ and SusCD_NoPlug_ structures, a lopsided protrusion on the extra- cellular side of the β-barrel cradles the distal part of SusD (**Fig 2A-H**). This protrusion is formed by EL3 (residues 371-437), EL1 (residues 259-302), EL10 (residues 877-932), and EL9 (residues 775–854), with the N-terminus of SusD resting on EL7 (residues 640-687) and EL8 (residues 714-738; **Fig 3A-C).** The SusC β-barrels are similar across all three unliganded structures, with SusC from SusCD_NoPlug_ and SusCD_+Plug_ superposing on SusC from SusCD_Open_ with RMSDs of 1.9Å (over 698 Ca pairs) and 0.8Å (over 805 Ca pairs), respectively (**Fig 3D-F**). The most notable difference among the three SusC β-barrels is that the SusCD_Open_ structure lacks density for EL7 and EL8 (**Fig 2I-L**, **Fig 3F**). In the SusCD_NoPlug_ and SusCD_+Plug_ conformations, SusD residues 25-43 wrap around the side of the SusC barrel at the base of EL7 (**Fig 3C**). This region has been described in other *Bacteroides* SusC-like proteins as a hinge that allows the SusD-like protein to pivot open (20, 26).

**Figure 3.**
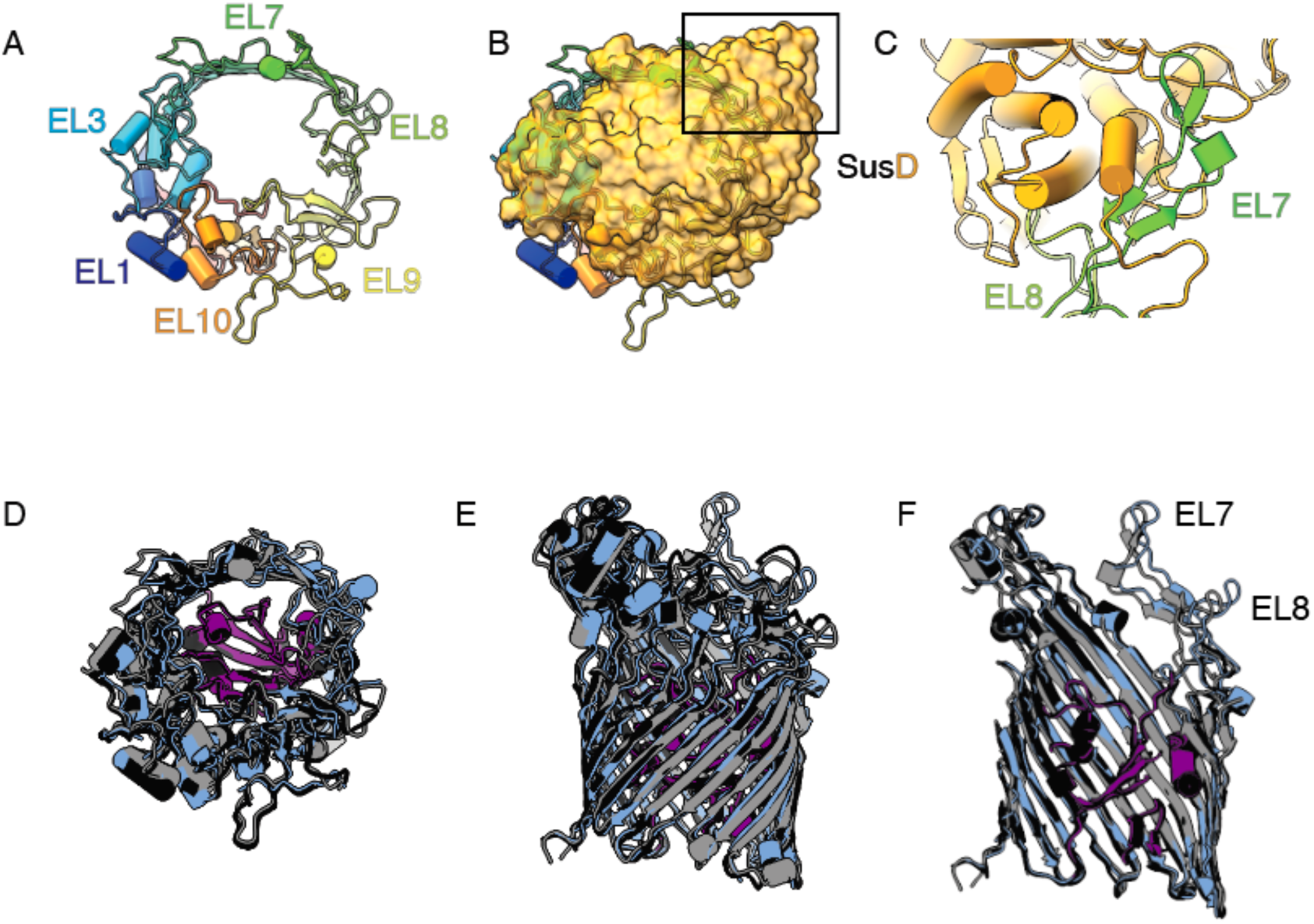
SusC extracellular loops interact with SusD. A. Top-down (extracellular) view of SusC with individual extracellular loops (ELs) that interact with SusD labeled. B. Similar view as in A with SusD shown in yellow space-filled model. Boxed region indicated where hinge loops EL7 and EL8 interact with SusD. C. Close-up view of the Nterminus of SusD that wraps around EL7 and EL8. D-F. Superposition of the SusCs from unliganded complexes with SusCD_+Plug_ in blue (barrel) and purple (plug), SusCD_NoPlug_ in gray and SusCD_Open_ in black. D is the top-down (extracellular) view similar as panel A, while the view in F is cropped in the vertical plane to show the plug domain.

### Maltoheptaose (M7) binding at two distinct sites stabilizes SusCD in a closed heterodimeric conformation

Previous work established that SusCD is required for *Bt* uptake of maltoheptaose (M7) and that recombinant SusD binds this carbohydrate with a Kd of ∼1mM (16, 17). To determine whether ligand binding alters the conformation of SusCD, we incubated SusCD with 10mM M7 for one hour on ice before vitrification using ultrathin carbon grids and collected images on a 300 kV Titan Krios equipped with a Gatan K3 direct electron detector and a Bioquantum energy filter (**Fig S5A,B**). 2D classification of vitrified SusCD_+M7_ particles have clear secondary structural features and there appear to be multiple views of the complex (**Fig S5C**). As observed for the unliganded particles, the M7-bound complex also appears heterodimeric in these averages.

Using single particle cryo-EM image processing methods, we determined the structure of SusCD bound to M7 (SusCD_+M7_) to a global resolution of 2.3 Å (**Fig 4A-D, Fig S5, Table S1**). The quality of the cryo-EM density map was sufficient to directly model SusC residues 107 through the C-terminus. As in the unliganded structures, density for the NTE was not observed. Likewise, the density for the lipoprotein SusD was sufficient to model the entire mature protein (residues 25-551), including a single acyl chain attached to Cys25 via a thioester linkage.

**Figure 4.**
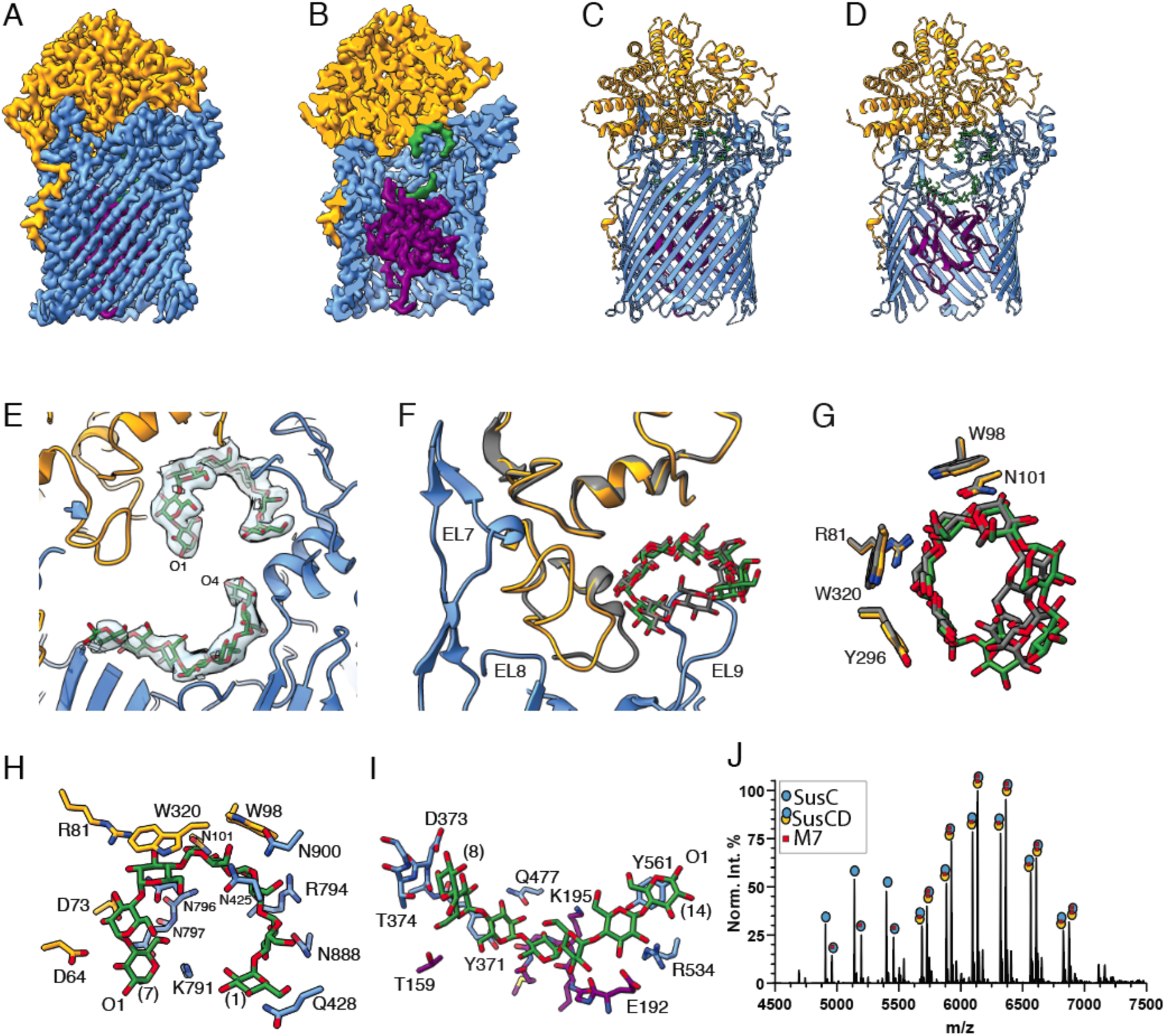
CryoEM structure of SusCD with maltoheptaose reveals a tightly closed complex. A-D. 2.3Å resolution electron density map of the maltoheptaose (M7)-bound SusCD complex with SusD in yellow, the SusC barrel in blue, the SusC plug in purple, and M7 in green. Panels B and D are clipped in the vertical plane to show the barrel interior. E. Close-up of the electron density for M7. O1 designates the reducing end of M7 at site one (top) and O4 designates the nonreducing end of M7 at site two (bottom). F. Close-up overlay of the SusD crystal structure with M7 (gray, PDB 3CK9) with the SusCD_+M7_ structure demonstrating the rearrangement of SusD loop T58-D73. G. Close-up overlay of the residues that coordinate M7 in the SusD crystal structure (gray, PDB 3CK9), and site one of the SusCD_+M7_ structure. H. Close-up view of site one with SusD in yellow, SusC in blue and M7 in green and red sticks. The M7 non-reducing and reducing ends are labeled as (1) and (7), respectively, and the reducing end O1 is labeled. I. Closeup view of site two with the SusC barrel in blue and the plug domain in purple, with M7 as green and red sticks. The M7 non-reducing and reducing ends are labeled as (8) and (14) respectively, and the reducing end O1 is labeled. For both panels H and I, residues within 3.5Å are displayed, as well as aromatic residues within ∼4Å. J. High-resolution native MS data of 3 μM SusCD with 5 mM M7.

Lipoproteins in *Bt* are likely triacylated, although the precise post-translational modification for SusD, or any lipoprotein in this organism, has not been defined (36).In the SusCD_+M7_, we observe ligand density at two sites: site 1, located at the SusC-SusD interface, and site 2, located deeper within the SusC β-barrel at the top of the plug (**Fig 4A-E**). The SusD T58-D73 loop shifts to accommodate M7 and interacts with SusC at EL7 and EL8 (**Fig 4F**). It does not appear that comparable conformational changes in SusD-like proteins upon ligand binding, or upon interaction with their cognate SusC partners have been reported for other structurally defined SusCD-like complexes (20, 24, 25). The changes in SusD do not appear to directly contribute to M7 binding, as the sugar-SusD contacts in the SusCD_+M7_ structure are identical to those observed in the SusD_+M7_ crystal structure (PDB 3CK9, **Fig 4G**) (16). The functional significance of this flexibility is not obvious from this structure. However, SusD G65, which is found within the flexible T58-D73 loop, is ∼10 Å from the reducing end of glucose (Glc14) of the ligand bound at site 2, potentially constraining the size of the ligand accommodated at this site.

At site 1, Glc1 and Glc2 at the non-reducing end of M7 interact exclusively with SusC residues Q428, R794, N888, and N900 (**Fig. 4H**). Toward the reducing end, Glc3–Glc7 are sandwiched between SusC residues N425, N900, N796, N797, and K791 and SusD residues W98, N101, W320, R81, D73, and D64. The reducing end O1 atom of Glc7 at site 1 is ∼7 Å from the non-reducing end O4 atom of Glc14 at site 2 (**Fig 4E**). This distance is greater than the length of a single α1,4-linked glucose (∼4.3 Å). We previously showed that SusCD preferentially takes up α1,4-linked glucans of 15-17 residues, which is consistent with the cryo-EM structure of the levan-targeting SusCD-like complex BT1763–BT1762 bound to a β2,6-linked fructooligosaccharide of 15 monosaccharides (∼2.5 kDa) (20, 37). The two M7 ligands observed in the SusCD_+M7_ structure likely capture the key interaction determinants for an exclusively α1,4-linked glucan, regardless of whether the transporter accommodates one longer MOS or two shorter MOS molecules.

The second M7-binding site in SusC is formed by the β-barrel wall and the top of the plug domain (**Fig 4I**). From the non-reducing (Glc8) end to the reducing end (Glc14), site 2 includes β-barrel residues D373, T374, Q477, Y371 R534, and Y561, as well as plug residues T159, E192, and K195. In addition, many of the Glc O2, O3 and O6 atoms, particularly towards the non-reducing end, interact with the peptide backbone, helping to lock this portion of the ligand in place.

We also used native MS to analyze the SusCD complex in the presence of M7, using 3mM SusCD complex after incubation with 5mM M7. This analysis found two predominant ion species for the SusCD complex and two main species for SusC alone (**Fig 4J**). For SusCD, we detected complexes bound to either one or two M7 molecules, with the two-ligand bound state as the predominant species at ∼55% of the sample. Notably, no unliganded SusCD was observed. In contrast, SusC was detected both with and without M7, consistent with SusD increasing M7 binding affinity. Approximately 32% of SusC signal corresponded to the M7- bound form, demonstrating that SusD is not strictly required for SusC to bind M7 *in vitro*. This finding aligns with our previously published data showing that a binding-deficient SusD (alanine substitution of W98, Y296 and W320) impairs, but does not eliminate, growth on maltooligosaccharides (38). Together, these data support a model in which M7 binds the SusCD complex at two distinct sites and in which ligand binding to SusC is not strictly dependent on prior loading by the SusD binding site.

### M7 binding stabilizes a closed SusCD conformation

Comparison of the SusCD_+M7_ structure with the unliganded SusCD_+Plug_ structure shows that SusC has a similar conformation in both states, with an RMSD of 1.0 Å over 858 Ca (**Fig 5A**). By contrast, SusC in the SusCD_+M7_ differs conformationally from SusC in the SusCD_NoPlug_ model. Superposition of SusC from SusCD_+M7_ with SusC from SusCD_NoPlug_ yields an RMSD of 2.0 Å over 755 Ca (**Fig 5B**). Comparison of the SusCD_+M7_ model with the SusCD_NoPlug_ reveals differences both at the SusC-D hinge region and at the distal closure point, where SusD contacts SusC at EL1. In the SusCD_+M7_ structure, these contact points are closer together: the distance between SusD T351 and SusC P284 (of EL1) is <5 Å in SusCD_+M7_, compared with 13 Å in unliganded SusCD_NoPlug_ and in the SusCD_+Plug_ structures. These differences suggest that MOS binding promotes a more “sealed” conformation of the complex, which may help initiate the transport cycle. The other pronounced difference is at the base of the hinge loops EL7 and EL8 in each structure, which are shifted toward the β-barrel lumen in the SusCD_NoPlug_ model relative to their conformation in SusCD_+M7_ (**Fig 5C**). The position of the EL7 and EL8 loops in SusCD_NoPlug_ would clash with the plug domain, particularly with the plug loop spanning residues 169-181. This suggests that displacement or loss of the plug is coupled to conformational changes in SusC.

**Figure 5.**
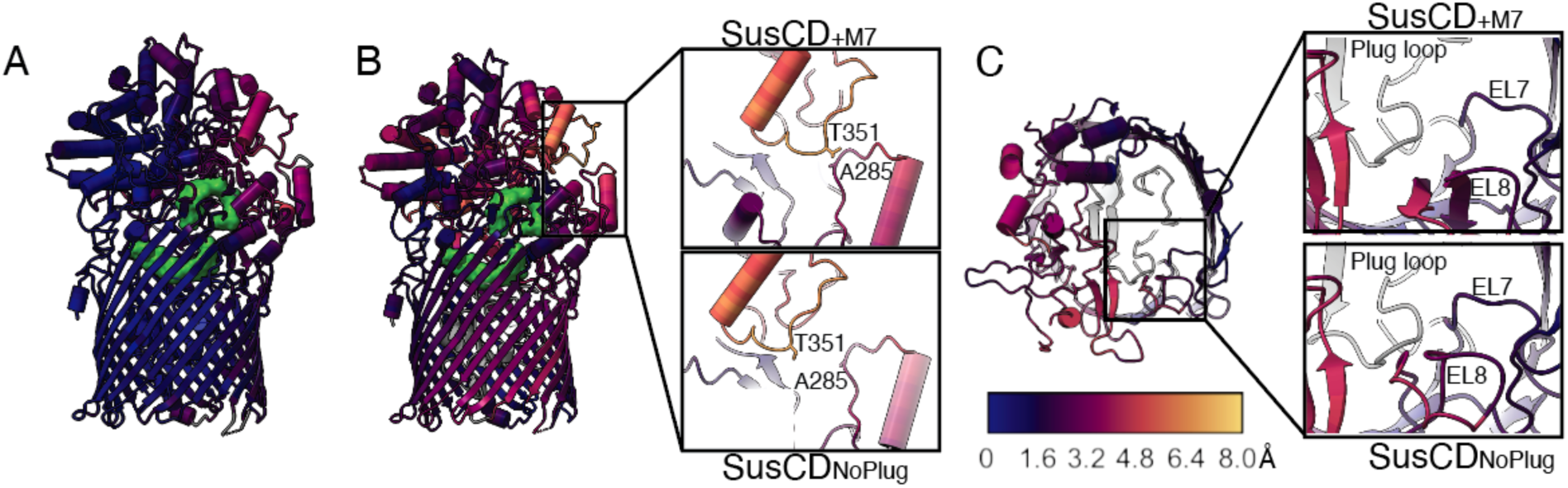
Comparison of the liganded and unliganded SusCD complexes demonstrate positional changes in both proteins. A. Average displacement of aligned residues between the SusCD_+M7_ complex (shown) versus the coordinates for SusCD_+Plug_ using the program ResiRuler. Coordinate distance key in A is shown in panel C. B. Average displacement of aligned residues between the SusCD_+M7_ complex (shown) versus the coordinates for SusCDNoPlug. Boxed region and inset shows the change in SusD closure over SusC for the most distal pivot point between the proteins. C. Average displacement of aligned residues between the SusCD+M7 complex (shown) versus the coordinates for SusCD_NoPlug_ as in panel A viewed from the top down (extracellular side). Insets demonstrate the positional change in EL7 and EL8 away from the plug loop (gray) in SusCD_+M7_ (top) versus SusCD_NoPlug_. The plug loop of SusCD_+M7_ (gray) is shown in SusCD_NoPlug_ inset.

Despite these differences in SusC EL7 and EL8 loops, we did not observe discrete changes in plug position or plug conformation in SusCD_Open_ and SusCD_+Plug_, the two unliganded complex conformations with density for the plug. We also did not detect positional changes in the C-terminal portion of the TonB box, previously defined as D105-V109, when comparing the unliganded structures with SusCD_+M7_ (20, 39). Although D105 and E106 of the DEVVV TonB box in SusC could be modeled in the unliganded SusCD structures, these two residues could not be confidently modeled in the SusCD_+M7_. However, the three valines of the TonB box adopt the same positions in all structures, regardless of whether M7 is bound. The inability to model residues D105 and E106 in SusCD_+M7_ may reflect increased local flexibility upon ligand binding. However, if M7 binding substantially altered TonB box presentation, we would expect more pronounced changes across the TonB box region as a whole.

## Discussion

In this study we determined structures of the unliganded and ligand-bound Bt SusCD complex. These structures reveal how the transporter may undergo conformational changes during substrate transport and highlight structural features that distinguish this system from other SusCD-like complexes. Our structural analysis supports the “pedal-bin” mechanism of transport proposed for other SusCD-like complexes, whereby lid closure of SusD on SusC precedes opening of the barrel on the periplasmic side to enable ligand transport (26).

We hypothesize that our structures capture several key changes in SusCD that occur during the transport cycle (**Fig 6A**). Prior to transport and as suggested by the SusCD_Open_ structure and 3DVA, SusD flexibly samples a wide range of conformations in relation to SusC. This is further supported by the lack of density for SusC EL7 and EL8 in the SusCD_Open_ complex, consistent with a model in which these loops become ordered when SusD is stably closed over SusC (**Figs 2,3**). While SusD can close over SusC in the absence of ligand, the complex doesn’t seem to lock shut without ligand (**Fig 5B, Movie S3**). Addition of ligand showed two distinct M7 binding sites: site 1, located at the SusC-SusD interface, and site 2, within the SusC β-barrel at the top of the plug. Binding of these two M7 molecules stabilizes the SusCD closed conformation which likely represents an intermediate state within the transport cycle.

**Figure 6:**
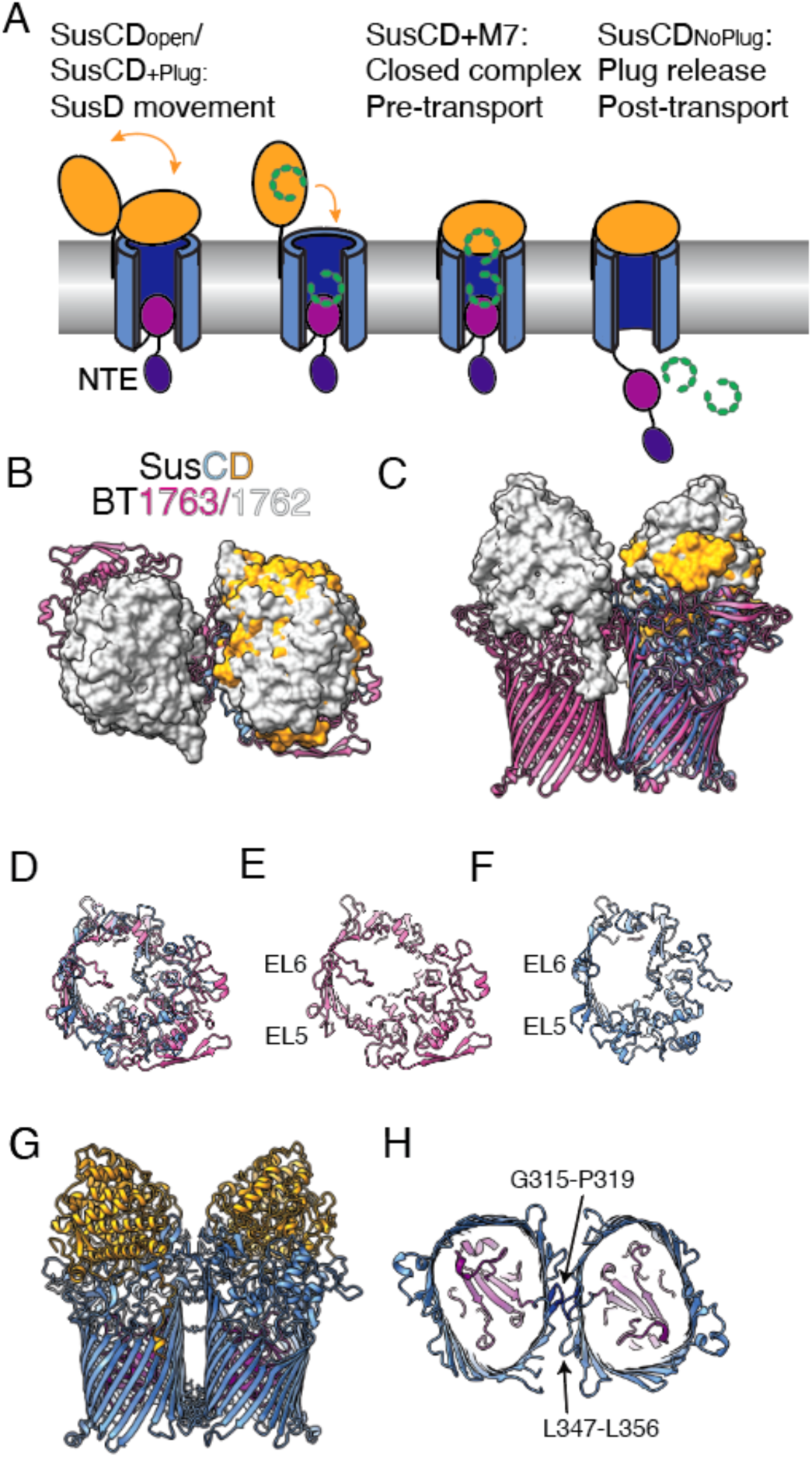
Working model of MOS transport by SusCD and structural basis for the homodimeric structure. A. The unliganded and M7 structures of SusCD likely reflect the conformations of the complex during the transport cycle. B. Overlay of the heterotetramer of BT1763-1762 (PDB 8AA3) with the heterodimer of SusCD viewed from the top down (extracellular side) C. Side-on view of the same overlay as in panel B. D. Overlay of one monomer of BT1763 (pink) and SusC (blue) viewed from the top-down (extracellular side) E, F. BT1763 and SusC as in panel D with the dimer loops of BT1763 (EL5 and EL6) labeled. G. Dimer of SusCD_+M7_ created by superposition onto BT1763-1762 (PDB 8AA3). Residues that clash between the SusC monomers as calculated in ChimeraX are displayed. H. Bottom-up (periplasmic) view of the SusC dimer in panel G, displaying the loops G315- P319 (dark blue) and L347-L356 that overlap and prevent SusC dimerization.

It is notable that we observe unliganded SusCD particles with and without density for the plug domain, as it has been hypothesized that the plug dissociates completely from the barrel of SusC-like proteins during transport (26). Our data supports this model. Although it is possible that the SusCD_NoPlug_ model represents a small subset of particles in which the plug was proteolyzed, we think it is more likely that this model represents a complex after ligand transit. Given the variable size and shape of MOS and similarly complex ligands transported by SusCD-like complexes, it seems likely that the plug must be ejected rather than squeezed or shifted within the β-barrel as hypothesized for Proteobacterial TBDTs that function without a lid-like protein (18, 19).

TBDTs are energized by β-strand pairing between the TonB box sequence which precedes the plug domain and the C-terminal domain of the inner membrane TonB complex. In previous work, we showed that deletion of the SusC TonB box eliminates growth on maltoheptaose and starch, supporting its importance in pairing with TonB (39). However, it remains possible that the NTE of SusC also contributes to TonB engagement and helps initiate movement of the TonB box, and perhaps the plug, during the transport cycle. A precedent for an important role of the NTE comes from the levan-targeting SusCD complex BT1763-BT1762. In the structure of this complex bound to fructooligosaccharides, the BT1763 TonB box was ejected from the transporter. However, deletion of the BT1763 TonB box only moderately influenced growth, where the deletion of the BT1763 NTE eliminated growth on fructooligosaccharides (20). Flexibility of the NTE, which is supported by that fact that we only observe NTE-like density through 3DVA, may allow this domain to find its cognate TonB in the periplasm. *Bacteroides* have multiple TonB proteins, and the NTE may impart some level of specificity to the TBDT-TonB interaction (40, 41).

Our structure of SusCD also reveals key differences from other SusCD-like complexes studied in *Bacteroides* and *Porphyromonas*, most notably in the stoichiometry of the complexes. Where our data demonstrates that SusCD is a heterodimer, other SusCD-like complexes are heterotetramers containing two copies each of SusC and SusD and some have been isolated with additional lipoprotein components (20, 24–27). We compared the structure of the well-studied levan-targeting SusCD-like complex BT1763-1762 with ligand (PDB 8AA3) to that of SusCD_+M7_ (**Fig 6B,C**) (27). SusC and BT1763 overlay with an RMSD of 0.9 Å for 554 Ca atoms (6.7Å over 824 Ca). For BT1763 and all other structurally characterized SusCD-like complexes, dimerization is mediated between the two SusC-like proteins, allowing the SusD lids to function independently. A common feature in these SusC-like dimers is the swapping of EL5 between the monomers, which might help to stabilize the dimer interface. In heterotetrameric SusCD-like structures, EL5 generally lacks defined secondary structure, whereas EL5 of SusC in our SusCD structures is similar in length but contains a short β-hairpin, suggesting a more rigid structure that would impede SusC-SusC interactions (**Fig 6D-F**).

Because our data do not exclude the possibility that SusCD forms a heterotetramer *in vivo*, we tried to create a *Bt* SusC dimer in two ways. First, we tried AlphaFold3, requesting a complex of SusC_2_D_2_ or SusC_2_ (42). These multimer predictions did not recapitulate the dimeric assembly (across EL5) that is conserved in all other SusCD-like complexes and returned low confidence models for SusC_2_D_2_ (ipTM = 0.43 and pTM=0.53) and SusC_2_ (ipTM=0.23, pTM=0.54), and the interfaces excluded EL5.

We then created a dimer of *Bt* SusCD by superimposing two SusC monomers from the SusCD_+M7_ structure onto BT1763-1762 with ligand (PDB 8AA3) (27). While this places EL5 of SusC in the dimer interface, there is significant clashing between adjacent SusC subunits at the base (periplasmic side) of the barrel, with a total of 251 clashes among 27 residues (**Fig 6G,H**). Significant overlap occurs between β-hairpin loops Gly315-Pro319 and L347-L356 in SusC while the same region in BT1763, loops defined by S316-G319 and L352-L357, adopt a different conformation (**Fig 6H**). We also created dimers by overlaying our SusC monomers onto four other determined SusCD-like structures, but the results were the same (20, 24, 27, 34). Therefore, both the conformation of the loops at the β-barrel base (periplasmic side) and the structure of EL5 contribute to the lack of *Bt* SusC dimerization. This provides a structural rationale for our observations that SusCD assembles as a heterodimer and has implications for how the Sus lipoproteins that support MOS uptake, such as SusE and SusG, can interact with SusCD.

To determine if our observations of the *Bt* SusC structure are likely conserved across other *Bacteroides* Sus PUL, we compared the sequences of 18 SusC homologs from type species of *Bacteroides* that have been validated to grow on starch (**Fig S6**) (32). SusC homologs were identified by their localization within putative operons that include a homolog of SusG and/or SusE. Multisequence alignment of the SusCs reveals that the most highly conserved region is at the N-terminus, including the NTE, TonB box and the plug domain (**Fig S7**). Many of the MOS binding residues are well conserved among nearest neighbors, with some variation seen in more distant homologs, which might reflect differences in size and shape of a-glucans transported. The most obvious divergent regions across SusC homologs occurs at extracellular loops, namely EL1, EL3, EL4, EL5, and EL9. This may simply reflect variation among homologs despite conserved assembly and function or differences in association with surface glycan binding proteins or SusG homologs. Finally, the intracellular loops that clash in our modeled SusC dimer are highly conserved. We repeated our AlphaFold3 search with the SusCD homologs from *Bacteroides ovatus* and *Bacteroides fragilis*, and these also favored a heterodimeric complex of SusC and SusD. While not conclusive, this does suggest that Sus complexes across *Bacteroides* are likely to have a monomeric SusC core.

While dimers of SusC-like proteins are retained under various detergent extraction protocols, larger complexes can be isolated using a single dodecylmaltoside extraction (27). These complexes, called “utilisomes”, capture a SusCD-like heterotetramer core and the surface glycan-binding lipoproteins and glycoside hydrolases encoded within the PUL. Thus, it has been hypothesized that complete utilisomes are assembled as functional units *in vivo* and deployed to degrade and capture carbohydrate nutrition. Our previous work has suggested that SusE and SusG, a starch-targeting surface glycan-binding protein and amylase respectively, dissociate from SusCD and may interact in a more dynamic fashion (29). In support of this, the structure of SusC has fewer potential docking points for additional lipoproteins, an observation first noted by White and colleagues in comparing the utilisome complexes for the capture of levan or dextran with a predicted *Bt* SusC structure (27). This can be seen in **Fig 6B,D-F**, which shows that the extracellular loops of BT1763 are more elaborate, while SusC is more compact. In summary, these differences in *Bt* SusCD compared to BT1763-1762 and other SusCD-like complexes, support our previous work on Sus function, and suggest that Sus, once considered a model for studying PUL-encoded carbohydrate uptake, is more of the exception than the rule.

Gut *Bacteroides* have evolved to forage on the vast array of carbohydrates that transit the gut by deploying many, and sometimes dozens of, highly specific SusCD-like transporters. This capacity enables gut microbes to adapt to the shifting nutrient microenvironment of the gut as the availability of specific carbohydrates changes. Our work shows that although key features of the SusCD-like transport are conserved between different PULs, these complexes are not monolithic and often have important structural differences. Understanding how these systems have diverged in form, and how this impacts function, will reveal the fundamental molecular principles guiding nutrient uptake in an abundant class of transporters within the *Bacteroides*.

## Experimental Procedures

### Construction of Bt susD-6xhis strain

The His-tagged SusD strain of *Bt* was created in the VPI 5482 *tdk-* background through methods previously described (16). Briefly, pExchange plasmid was used for homologous recombination of the *susD-6xhis* allele into Bt Δ*susD* via conjugation with *E. coli* S17 λ pir strain (16). Primers for cloning are listed in **Table S2**. Transformants were selected on BHI-blood plates supplemented with gentamycin (200mg/ml) and erythromycin (25mg/ml). Positive transformants were then counter selected by plating on BHI-blood plates supplemented with fluorodeoxyuridine (FUdR, 200mg/ml) to select for isolates that lost the vector backbone. Isolates were then selected for loss of erythromycin resistance and Sanger sequencing was used to confirm the *susD-6xhis* allele insertion at its native chromosomal location in Bt *tdk-*.

### Expression and Purification of SusCD Complex

*Bt* cultures were grown in a 37°C Coy anaerobic chamber (5% H_2_/10% CO_2_/85% N_2_). *Bt susD- 6xhis* was inoculated from a freezer stock into 10ml of TYG medium and grown for 18 hours in the anaerobic chamber to late stationary phase. This culture was used to inoculate 4L of *Bacteroides* minimal media with 5g/L maltose as the sole carbon source to upregulate *sus* (43). Cells were grown for 24 hours before harvesting via centrifugation (20K xg for 20 minutes).

The purification procedure for *Bt* SusCD was adapted from methods used for the purification of other SusCD-like complexes (26). Bt *susD-6xhis* cells were resuspended in 40mL of buffer A (25mM NaH_2_PO_4_, 500mM NaCl, 20mM Imidazole, pH7.5) supplemented with lysozyme, DNaseI, and one COMPLETE EDTA-free protease inhibitor tablet (Roche) and lysed by French press via two passes at 1200 psi. The lysate was centrifuged at 10K xg for 10 minutes to remove unbroken cells, and the supernatant was ultracentrifuged at 4°C for 1 hour at 41.5k RPM (220,780 xg) in a Thermo Scientific Sorvall WX Ultra Series centrifuge with a Fiberlite F50L-8x39mm rotor. Pellets containing total membrane were resuspended in 50mL buffer A with one COMPLETE EDTA free protease inhibitor tablet. Sodium lauryl sarcosine was added to a final concentration of 0.5% and stirred for 1 hour at room temperature to solubilize inner membrane proteins. This solution was ultracentrifuged again at 41.5K RPM (220,780 xg) for 30 minutes to pellet the outer membranes. The pellet was resuspended in buffer A with protease inhibitor, and β*-*octyl-glucoside was added to a final concentration of 2%. The solution was stirred overnight (16 hours) at 4°C; in testing different lengths of time for solubilization, overnight incubation did not have deleterious effects on the sample. A final ultracentrifugation step of 41.5k RPM (220,780 xg) was used to pellet insoluble material, and the supernatant containing solubilized outer membrane proteins was applied to a 5mL His-trap fast flow column (Cytiva) equilibrated with buffer A containing 0.15% dodecyl maltoside (DDM). The column was washed with 45 mL buffer A with 0.15% DDM and proteins were eluted with buffer B (25mM NaH_2_PO_4_, 500mM NaCl, 300mM Imidazole, pH7.5) containing 0.15% DDM. Samples containing SusCD were pooled via SDS-PAGE analysis and concentrated to <1mL via an Amplicon concentrator with a 10kDa MWCO. Concentrated sample was loaded onto a 10/300 increase Superdex 200 size exclusion column equilibrated in SEC buffer (20mM HEPES, 100mM NaCl, 0.03% DDM w/v, pH 7.0), and fractions containing SusCD were pooled based on SDS-PAGE analysis.

### Native MS of SusCD Complex

SusCD was prepared as described above with the exception that the SEC buffer (20mM HEPES, 100mM NaCl, 0.03% DDM w/v, pH 7.0) was modified to contain slightly over two times the critical micelle concentration of octyl-glucopyranoside detergent (OG), 0.8% w/v (26mM), in water. This detergent substitution was made due to issues encountered with liberating protein complex from DDM micelles during ionization, however we do not believe this significantly impacts complex behavior or ligand binding as the native MS data aligns with our finding via cryoEM.

Purified stock solutions of SusCD at 10-15mM were solvent exchanged into 200mM ammonium acetate with 0.8% OG using Amicon Ultra 0.5 ml centrifugal filters (MilliporeSigma) with a 10 kDa molecular weight cutoff. Three consecutive washing steps were performed for each sample. A stock solution of 100mM of maltoheptaose (M7) was prepared in 200mM ammonium acetate with 0.8% OG. To determine the oligomeric state, SusCD was diluted to 3µM for MS analysis. For M7 binding analysis, 3µM SusCD was incubated with 5mM M7 for 2 hours at 4°C. After incubation, samples were solvent exchanged again with 200mM ammonium acetate with 0.8% OG to remove any unbound M7.

High-resolution native MS analysis was performed on a Q Exactive Orbitrap Ultra High Mass Range (QE-UHMR) Hybrid Quadrupole-Orbitrap Mass Spectrometer (Thermo Scientific, San Jose, CA). ∼5mL of sample was introduced via nano electrospray ionization (nESI) in positive ion mode using gold-coated borosilicate capillaries needles prepared in-house. All data was collected by scanning 1,000-15,000 m/z with a resolution of 12,000 (at m/z 400). A capillary voltage of 1.1 kV was applied, while the capillary temperature was set to 250 °C. In-source collision-induced dissociation of 50V and higher-energy collisional dissociation (HCD) of 50V (in the ion-routing multipole) was employed to liberate protein ions from detergent micelles. High *m*/*z* detector and high *m*/*z* transfer optics were used, and the trapping gas pressure was set to five. Transient times were set at 100 ms and one minute (∼85 scans) of data collection was averaged for the presented mass spectra. Four measurements for each sample were performed. All acquired mass spectra were deconvoluted and analyzed using UniDec software (44).

### Cryo-EM sample preparation and data collection

Cryo-EM data collection parameters are summarized in **Table S1**. For the unliganded SusCD complex, 3.0 µl of SusCD (0.54 mg/mL) in SEC buffer (20mM HEPES, 100mM NaCl, 0.03% DDM w/v, pH 7.0) was applied to a glow discharged copper Quantifoil grid R1.2/1.3 with Ultrathin Carbon on a 300-mesh size. For the M7 complex, SusCD (0.95 mg/ml) in SEC buffer (20mM HEPES, 100mM NaCl, 0.03% DDM w/v, pH 7.0) was incubated with 10mM M7 for one hour on ice before 3 µl of sample was applied two times to a glow discharged copper Quantifoil grid R1.2/1.3 with Ultrathin Carbon on a 300-mesh size. Each application to the grid was incubated for 60s at room temperature and excess liquid was removed between sample applications by manual blotting at the edge of the grid. Grids were vitrified by plunging into liquid ethane slurry using a Mark IV Vitrobot (Thermo Fisher) at 4°C and 100% humidity using a blot force of 0 and blot time of 3s.

Micrographs were collected on a Titan Krios G3 (Thermo Fisher) operated at 300 kV. The images were collected with a K3 Summit direct electron detector (Gatan) equipped with a BioQuantum energy filter operating in counting mode. Collection for unliganded SusCD was done at a nominal magnification of 81,000X, corresponding to a pixel size of 1.08 Å and for M7 bound SusCD at a nominal magnification of 105,000 corresponding to a pixel size of 0.84 Å. The energy slit of the BioQuantum was set at a width of 20 eV. The total dose was 60 e/Å2 for unliganded collection and 55 e/Å2 for the M7 bound complex collection, fractionated over 60 frames. Data were collected using Serial EM software (version 4.0) with a nominal defocus range set from −1.5 to −2.5 μm.

### Cryo-EM data processing

Cryo-EM data processing and statistics are summarized in **Table S1**. CryoSPARC v. 4.0+ was used for all data processing (45). 4,549 movies of unliganded SusCD were aligned using patch motion correction and contrast transfer function (CTF) estimation was done using patch contrast transfer function. Particles were picked using Blob Picker using a diameter of 80-200 Å (or 50-200 Å for SusCD_Open_) and extracted with a box size of 360 pixels (384.2 Å).

Data processing and final map statistics for the unliganded data set are summarized in **Fig S2** and **Fig S3**, and the variability analysis (3DVA) workflow is in **Fig S4**. The unliganded SusCD micrographs were processed via two different workflows in which all three reconstructions (SusCD_+Plug_, SusCD_NoPlug_ and SusCD_Open_) were derived from both processing workflows as shown in **Fig S2**. All the maps were overall similar. The manual curation of images (right side **Fig S2**) eliminated those with excessive ice, and resulted in more complete maps of SusD with better local resolution throughout. The map used for model refinement is indicated in bold in **Fig S2**. The final resolutions for each unliganded SusCD map are 3.2 Å for the SusCD_Open_ map, 3.3 Å, for the SusCD_+Plug_, and 3.1 Å for the SusCD_NoPlug_ map. Gold standard Fourier shell correlation (GS-FSC), per-particle viewing direction distribution heat maps and local resolution is displayed in **Fig S3**. CryoSPARC 3DVA analysis was also used on the SusCD_Open_ particle stack (344,141 particles) that led to a map of SusCD_Open_ that resolved SusD in an open position (**Fig S4**). For 3DVA analysis, the SusCD_Open_ particle stack was processed using a 15Å filter resolution, then clustered into a discrete particle stack of particles found in component 0 (of three components). These particles were used to determine twenty 3D volumes. The volume with the best resolved SusD density in the open position was chosen as an initial model for non-uniform refinement using a particle stack (146,204 particles) that combined particles from other 3D classes with density for SusD in an open position. This SusCD_Open_ map reached a global resolution of 4.4Å.

For the SusCD_+M7_ data set, all data processing was performed in CryoSPARC v. 4.0+ (45). 3,410 movies were captured and aligned using patch motion correction and contrast transfer function (CTF) estimation was done using patch contrast transfer function. Particles were picked using Blob Picker using a diameter of 80-200 Å and extracted with a box size of 360 pixels (384.2 Å). Data processing and final map statistics for the liganded data set are shown in **Fig S5**, including Gold standard Fourier shell correlation (GS-FSC), per-particle viewing direction distribution heat maps and local resolution. The SusCD_+M7_ map achieved a 2.3 Å global resolution.

### Model building and visualization

Initial modeling of the SusCD complexes was done using rigid body docking in Phenix (v. 1.21.2) (46). A SusCD model was generated by Alphafold3 and docked into the CryoSPARC sharpened map densities using ChimeraX v1.10.1 (47). Pixel array spacing for the SusCD_+M7_ and the SusCD_Open_ map was optimized using Phenix.magref. Phenix real space refinement was used to refine the structure. Manual model building in Coot (v. 0.9.8.83) was used to adjust residue density fit and build in additional residues and molecules including M7 in the SusCD_+M7_ structure (48). Structures were visualized using ChimeraX v1.8 and ChimeraX v1.10.1 (47).

### Bacteroides SusC homolog analysis

As described, 18 SusC homologs were selected from predicted Sus loci in *Bacteroides* type strains that grow on starch and α-glucans as reported in Pudlo et al., 2022 (32). Sus loci were identified by inclusion of SusD, SusE and SusG encoded proteins with homology to those from *Bt* or *Bacteroides ovatus*, which have confirmed function. SusC proteins shared 32 – 83% identity. A multisequence alignment was created via CLUSTALW in MegAlignPro (DNAStar), and a neighbor-joining tree was constructed (49, 50). Gene sequences include *Phocaeicola massiliensis* HMPREF1534_02829, *Bacteroides thetaiotaomicron* BT_3702, *Phocaeicola plebius* BACPLE_02370, *Bacteroides intestinalis* BACINT_03131, *Bacteroides salyersiae* HMPREF1532_02495, *Bacteroides oleiciplenus* WP_009132665, *Bacteroides ovatus* Bovatus_03807, *Bacteroides xylanisolvens* BXY_47640, *Bacteroides cellulosyliticus* BcellWH2_01291 (sequence labeled 1), *Bacteroides fluxus* HMPREF9446_00914, *Bacteroides stercoris* BACSTE_01657, *Phocaeicola dorei* WP_008674305, *Phocaeicola vulgatus* BVU_1380, *Bacteroides finegoldii* BACFIN_07563, *Bacteroides cellulosyliticus* BcellWH2_02327 (sequenced labeled 2), *Bacteroides eggerthii* BACEGG_01900, *Bacteroides nordii* HMPREF1068_03059, *Bacteroides fragilis* BF3146, *Bacteroides uniformis* BACUNI_01209.

## Figure generation

Figures were schematized in Inkscape, Adobe Illustrator, Powerpoint, Chimera and ChimeraX (47).

## Data Availability

The SusCD_Open_, SusCD_+Plug_, SusCD_NoPlug_ and SusCD_M7_ EM density maps and models are deposited with the Electron Microscopy Data Bank (EMDB) and the RCSB Protein Data Bank (PDB) and will be released upon publication. Particle stacks are deposited in the Electron Microscopy Public Image Archive (EMPIAR) and will be released upon publication. EM density map for the SusCD_3DVA_ is deposited in EMDB and will be released upon publication.

## Code Availability

The UniDec software is available at https://github.com/michaelmarty/UniDec/. The ResiRuler software and documentation is available at https://github.com/tbaker67/ResiRuler/.

## Supporting information

Author Contributions

## Acknowledgements and Funding

The U-M Cryo-EM facility is supported by the UM Biosciences Initiative, the Arnold and Mabel Beckman Foundation, and the Life Sciences Institute. The U-M BioMS facility is supported by the UM Biosciences Initiative. This work was supported by the National Institutes of Health (NIH) S10OD030275 (M.D.O.), GM118475 (N.M.K), GM149374 (B.R.) and GM138620 (B.R.), F32AI186525 (W.S.) and an NSF-GRFP (N.R.F.) A University of Michigan Pandemic Research Recovery grant (N.M.K.) also supported this work. We thank Ashleigh Raczkowski, Vinson Lam, Alexandrea Rizo, Erica Murbach and Chris Lilienthal for cryo-EM facility support. We thank members of the Koropatkin and Ohi labs, and Tobias Giessen for helpful discussions.

## Author contributions

M.C.C., M.D.O. and N.M.K conceptualization;

M.C.C., N.R.F., W.S., J.R.R., B.R., M.D.O. and N.M.K. methodology and formal analysis;

M.C.C, N.R.F. and N.M.K. writing–original draft;

M.C.C, N.R.F., W.S., B.R., M.D.O, and N.M.K writing–review and editing;

B.R., M.D.O, and N.M.K. resources;

M.C.C, N.R.F., B.R., M.D.O and N.M.K. visualization;

B.R., M.D.O. and N.M.K. supervision and funding acquisition.

## References

1. Andreas J. Bäumler, and Vanessa Sperandio (2016) Interactions between the microbiota and pathogenic bacteria in the gut. Nature. 10.1038/nature18849

2. Wu, H.-J., and Wu, E. (2012) The role of gut microbiota in immune homeostasis and autoimmunity. Gut Microbes 10.4161/gmic.19320

3. Ding, R, Goh, W., Wu, R., Yue, X., Luo, X., Khine, W., Wu, J., and Lee, Y. (2019) Revisit gut microbiota and its impact on human health and disease. J. Food Drug Anal. 27, 623– 631

4. Makki, K., Deehan, E. C., Walter, J., and Bäckhed, F. (2018) The impact of dietary fiber on gut microbiota in host health and disease. Cell Host Microbe. 23, 705–715

5. Mukhopadhya, I., and Louis, P. (2025) Gut microbiota-derived short-chain fatty acids and their role in human health and disease. Nat. Rev. Microbiol. 23, 635–651

6. Zafar, H., and Saier, M. H. (2021) Gut Bacteroides species in health and disease. Gut Microbes. 13, 1–20

7. Sonnenburg, J. L., Xu, J., Leip, D. D., Chen, C.-H., Westover, B. P., Weatherford, J., Buhler, J. D., and Gordon, J. I. (2005) Glycan foraging in vivo by an intestine-adapted bacterial symbiont. Science (1979). **307**, 1952–1955

8. Martens, E. C., Chiang, H. C., and Gordon, J. I. (2008) Mucosal glycan foraging enhances fitness and transmission of a saccharolytic human gut bacterial symbiont. Cell Host Microbe. 4, 447–457

9. Grondin, J. M., Tamura, K., Déjean, G., Abbott, D. W., and Brumer, H. (2017) Polysaccharide Utilization Loci: Fueling microbial communities. J. Bacteriol. 199(15), e00860–16

10. Feng, J., Qian, Y., Zhou, Z., Ertmer, S., Vivas, E. I., Lan, F., Hamilton, J. J., Rey, F. E., Anantharaman, K., and Venturelli, O. S. (2022) Polysaccharide utilization loci in Bacteroides determine population fitness and community-level interactions. Cell Host Microbe. 30, 200–215.e12

11. McNulty, N. P., Wu, M., Erickson, A. R., Pan, C., Erickson, B. K., Martens, E. C., Pudlo, N. A., Muegge, B. D., Henrissat, B., Hettich, R. L., and Gordon, J. I. (2013) Effects of diet on resource utilization by a model human gut microbiota containing Bacteroides cellulosilyticus WH2, a symbiont with an extensive glycobiome. PLoS Biol. 10.1371/journal.pbio.1001637

12. Anderson, K. L., and Salyers, A. A. (1989) Biochemical evidence that starch breakdown by Bacteroides thetaiotaomicron involves outer membrane starch-binding sites and periplasmic starch-degrading enzymes. J. Bacteriol. 171, 3192–3198

13. Shipman, J. A., Cho, K. H., Siegel, H. A., and Salyers, A. A. (1999) Physiological characterization of SusG, an outer membrane protein essential for starch utilization by Bacteroides thetaiotaomicron. J. Bacteriol. 181, 7206–7211

14. Cameron, E. A., Maynard, M. A., Smith, C. J., Smith, T. J., Koropatkin, N. M., and Martens, E. C. (2012) Multidomain carbohydrate-binding proteins involved in Bacteroides thetaiotaomicron starch metabolism. Journal of Biological Chemistry. 287, 34614–34625

15. Shipman, J. A., Berleman, J. E., and Salyers, A. A. (2000) Characterization of four outer membrane proteins involved in binding starch to the cell surface of Bacteroides thetaiotaomicron. J. Bacteriol. 182, 5365–5372

16. Koropatkin, N. M., Martens, E. C., Gordon, J. I., and Smith, T. J. (2008) Starch catabolism by a prominent human gut symbiont is directed by the recognition of amylose helices. Structure. 16, 1105–1115

17. Reeves, A. R., D’elia, J. N., Frias, J., and Salyers, A. A. (1996) A Bacteroides thetaiotaomicron outer membrane protein that is essential for utilization of maltooligosaccharides and starch. J. Bacteriol. 178, 823–830

18. Noinaj, N., Guillier, M., Barnard, T. J., and Buchanan, S. K. (2010) TonB-dependent transporters: Regulation, structure, and function. Annu. Rev. Microbiol. 64, 43–60

19. Silale, A., and Van Den Berg, B. (2026) TonB-dependent transport across the bacterial outer membrane. Annual Review of Microbiology *Downloaded from* www.annualreviews.org. *Guest*. 48, 32

20. Gray, D. A., White, J. B. R., Oluwole, A. O., Rath, P., Glenwright, A. J., Mazur, A., Zahn, M., Baslé, A., Morland, C., Evans, S. L., Cartmell, A., Robinson, C. V., Hiller, S., Ranson, N. A., Bolam, D. N., and van den Berg, B. (2021) Insights into SusCD-mediated glycan import by a prominent gut symbiont. Nat. Commun. 10.1038/s41467-020-20285-y

21. D’elia, J. N., and Salyers, A. A. (1996) Contribution of a neopullulanase, a pullulanase, and an alpha-glucosidase to growth of Bacteroides thetaiotaomicron on starch. J. Bacteriol. 178, 7173–7179

22. D’elia, J. N., and Salyers, A. A. (1996) Effect of regulatory protein levels on utilization of starch by Bacteroides thetaiotaomicron. J. Bacteriol. 178, 7180–7186

23. Martens, E. C., Koropatkin, N. M., Smith, T. J., and Gordon, J. I. (2009) Complex glycan catabolism by the human gut microbiota: The Bacteroidetes sus-like paradigm. Journal of Biological Chemistry. 284, 24673–24677

24. Madej, M., White, J. B. R., Nowakowska, Z., Rawson, S., Scavenius, C., Enghild, J. J., Bereta, G. P., Pothula, K., Kleinekathoefer, U., Baslé, A., Ranson, N. A., Potempa, J., and van den Berg, B. (2020) Structural and functional insights into oligopeptide acquisition by the RagAB transporter from Porphyromonas gingivalis. Nat. Microbiol. 5, 1016–1025

25. Tong, M., Xu, J., Li, W., Jiang, K., Yang, Y., Chen, Z., Jiao, X., Meng, X., Wang, M., Hong, J., Long, H., Liu, S. J., Lim, B., and Gao, X. (2024) A highly conserved SusCD transporter determines the import and species-specific antagonism of Bacteroides ubiquitin homologues. Nature Communications. 10.1038/s41467-024-53149-w

26. Glenwright, A. J., Pothula, K. R., Bhamidimarri, S. P., Chorev, D. S., Baslé, A., Firbank, S. J., Zheng, H., Robinson, C. V., Winterhalter, M., Kleinekathöfer, U., Bolam, D. N., and Van Den Berg, B. (2017) Structural basis for nutrient acquisition by dominant members of the human gut microbiota. Nature. 541, 407–411

27. White, J. B. R., Silale, A., Feasey, M., Heunis, T., Zhu, Y., Zheng, H., Gajbhiye, A., Firbank, S., Baslé, A., Trost, M., Bolam, D. N., van den Berg, B., and Ranson, N. A. (2023) Outer membrane utilisomes mediate glycan uptake in gut Bacteroidetes. Nature. 618, 583–589

28. Martens, E. C., Lowe, E. C., Chiang, H., Pudlo, N. A., Wu, M., McNulty, N. P., Abbott, D. W., Henrissat, B., Gilbert, H. J., Bolam, D. N., and Gordon, J. I. (2011) Recognition and degradation of plant cell wall polysaccharides by two human gut symbionts. PLoS Biol. 10.1371/journal.pbio.1001221

29. Tuson, H. H., Foley, M. H., Koropatkin, N. M., and Biteen, J. S. (2018) The starch utilization system assembles around stationary starch-binding proteins. Biophys. J. 115, 242–250

30. Karunatilaka, K. S., Cameron, E. A., Martens, E. C., Koropatkin, N. M., and Biteen, J. S. (2014) Superresolution imaging captures carbohydrate utilization dynamics in human gut symbionts. mBio. 10.1128/mBio.02172-14

31. Brown, H. A., DeVeaux, A. L., Juliano, B. R., Photenhauer, A. L., Boulinguiez, M., Bornschein, R. E., Wawrzak, Z., Ruotolo, B. T., Terrapon, N., and Koropatkin, N. M. (2023) BoGH13ASus from Bacteroides ovatus represents a novel α-amylase used for Bacteroides starch breakdown in the human gut. Cellular and Molecular Life Sciences. 10.1007/s00018-023-04812-w

32. Pudlo, N. A., Urs, K., Crawford, R., Pirani, A., Atherly, T., Jimenez, R., Terrapon, N., Henrissat, B., Peterson, D., Ziemer, C., Snitkin, E., and Martens, E. C. (2022) Phenotypic and genomic diversification in complex carbohydrate-degrading human gut bacteria. mSystems

33. Punjani, A., and Fleet, D. J. (2021) 3D variability analysis: Resolving continuous flexibility and discrete heterogeneity from single particle cryo-EM. J. Struct. Biol. 10.1016/j.jsb.2021.107702

34. Abellon-Ruiz, J., Jana, K., Silale, A., Frey, A. M., Baslé, A., Trost, M., Kleinekathöfer, U., and van den Berg, B. (2023) BtuB TonB-dependent transporters and BtuG surface lipoproteins form stable complexes for vitamin B12 uptake in gut Bacteroides. Nat. Commun. 10.1038/s41467-023-40427-2

35. Silale, A., Soo, Y. L., Mark, H., Motz, R. N., Baslé, A., Nolan, E. M., and van den Berg, B. (2026) Structural basis of iron piracy by human gut Bacteroides. PNAS Microbiology. 10.1101/2024.04.15.589501

36. Armbruster, K. M., Jiang, J., Sartorio, M. G., Scott, N. E., Peterson, J. M., Sexton, J. Z., Feldman, M. F., and Koropatkin, N. M. (2024) Identification and characterization of the lipoprotein N-acyltransferase in Bacteroides. Proc. Natl. Acad. Sci. U. S. A. 10.1073/pnas.2410909121

37. Foley, M. H., Martens, E. C., and Koropatkin, N. M. (2018) SusE facilitates starch uptake independent of starch binding in B. thetaiotaomicron. Mol. Microbiol. 108, 551–566

38. Cameron, E. A., Kwiatkowski, K. J., Lee, B. H., Hamaker, B. R., Koropatkin, N. M., and Martens, E. C. (2014) Multifunctional nutrient-binding proteins adapt human symbiotic bacteria for glycan competition in the gut by separately promoting enhanced sensing and catalysis. mBio. 10.1128/mBio.01441-14

39. Pollet, R. M., Foley, M. H., Kumar, S. S., Elmore, A., Jabara, N. T., Venkatesh, S., Pereira, G. V., Martens, E. C., and Koropatkin, N. M. (2023) Multiple TonB homologs are important for carbohydrate utilization by Bacteroides thetaiotaomicron. J. Bacteriol. 10.1101/2023.07.07.548152

40. Parker, A. C., Seals, N. L., Baccanale, C. L., and Rocha, E. R. (2022) Analysis of Six tonB gene homologs in Bacteroides fragilis revealed that tonB3 is essential for survival in experimental intestinal colonization and intra-abdominal infection. Infect. Immun.

41. Rebecca M. Pollet, Lauryn M. Martin, and Nicole M. Koropatkin (2021) TonB-dependent transporters in the Bacteroidetes: Unique domain structures and potential functions. Mol. Microbiol. 115, 490–501

42. Abramson, J., Adler, J., Dunger, J., Evans, R., Green, T., Pritzel, A., Ronneberger, O., Willmore, L., Ballard, A. J., Bambrick, J., Bodenstein, S. W., Evans, D. A., Hung, C. C., O’Neill, M., Reiman, D., Tunyasuvunakool, K., Wu, Z., Žemgulytė, A., Arvaniti, E., Beattie, C., Bertolli, O., Bridgland, A., Cherepanov, A., Congreve, M., Cowen-Rivers, A. I., Cowie, A., Figurnov, M., Fuchs, F. B., Gladman, H., Jain, R., Khan, Y. A., Low, C. M. R., Perlin, K., Potapenko, A., Savy, P., Singh, S., Stecula, A., Thillaisundaram, A., Tong, C., Yakneen, S., Zhong, E. D., Zielinski, M., Žídek, A., Bapst, V., Kohli, P., Jaderberg, M., Hassabis, D., and Jumper, J. M. (2024) Accurate structure prediction of biomolecular interactions with AlphaFold 3. Nature. 630, 493–500

43. Foley, M. H., Cockburn, D. W., and Koropatkin, N. M. (2016) The Sus operon: a model system for starch uptake by the human gut Bacteroidetes. Cellular and Molecular Life Sciences. 73, 2603–2617

44. Marty, M. T., Baldwin, A. J., Marklund, E. G., Hochberg, G. K. A., Benesch, J. L. P., and Robinson, C. V. (2015) Bayesian deconvolution of mass and ion mobility spectra: From binary interactions to polydisperse ensembles. Anal. Chem. 87, 4370–4376

45. Punjani, A., Rubinstein, J. L., Fleet, D. J., and Brubaker, M. A. (2017) CryoSPARC: Algorithms for rapid unsupervised cryo-EM structure determination. Nat. Methods. 14, 290–296

46. Afonine, P.V., Poon, B.K., Read, R.J., Sobolev, O.V., Terwilliger, T.C., Urzhumtsev, A., Adams, P.D. (2018) Real-space refinement in PHENIX for cryo-EM and crystallography. Acta Crystallogr D Struct Biol. 74, 531–544

47. Meng, E. C., Goddard, T. D., Pettersen, E. F., Couch, G. S., Pearson, Z. J., Morris, J. H., and Ferrin, T. E. (2023) UCSF ChimeraX: Tools for structure building and analysis. Protein Science. 10.1002/pro.4792

48. Emsley, P., and Cowtan, K. (2004) Coot: Model-building tools for molecular graphics. Acta Crystallogr. D Biol. Crystallogr. 60, 2126–2132

49. Thompson, J. D., Gibson, Toby. J., and Higgins, D. G. (2003) Multiple Sequence Alignment Using ClustalW and ClustalX. Curr. Protoc. Bioinformatics. 10.1002/0471250953.bi0203s00

50. Burland, T. G. DNASTAR’s Lasergene Sequence Analysis Software

