## Supplemental Figures for "Structural analyses of a heterodimeric SusCD complex captures intermediate states of maltooligosaccharide transport"

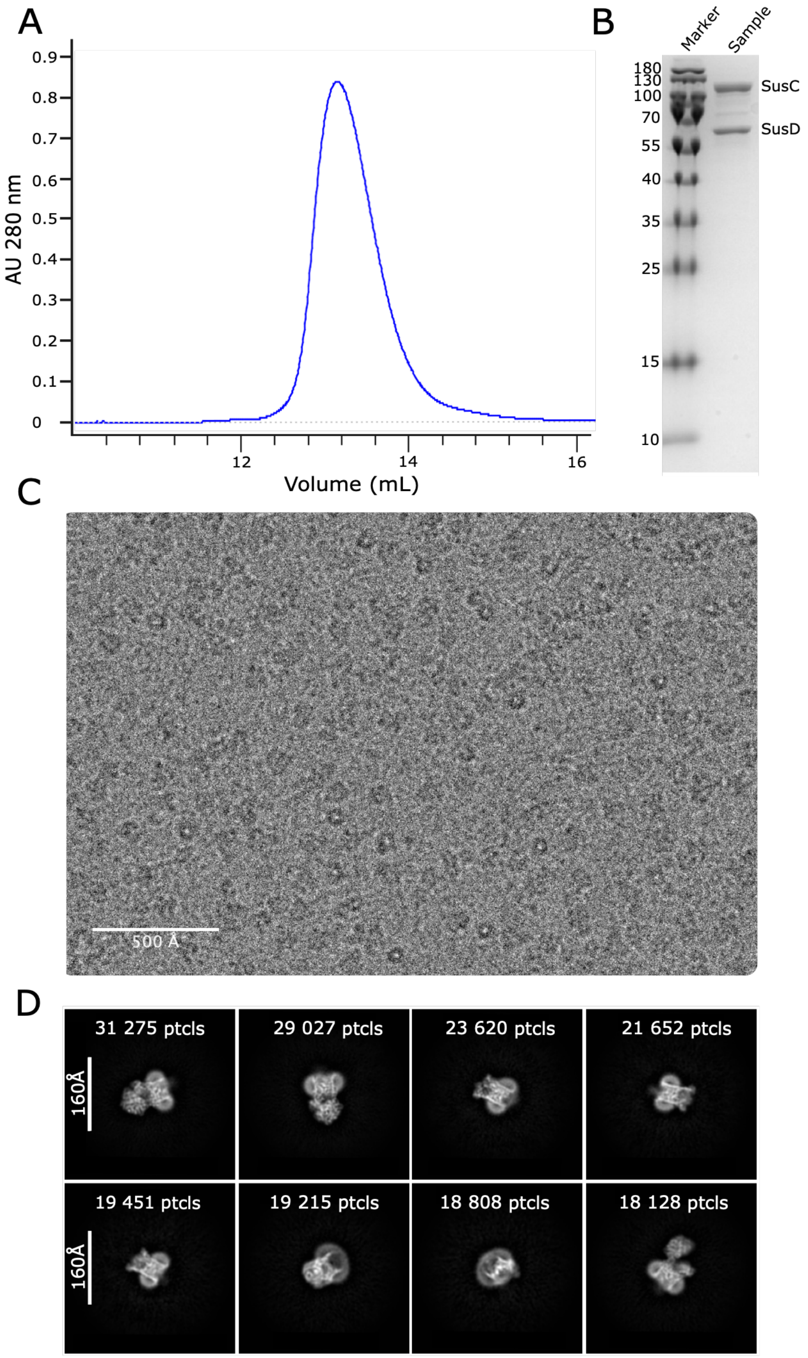


**Supplemental Figure 1. Purification and 2D classes for unliganded SusCD**. A. Superdex 200 size exclusion chromatography yielded a single peak at ~13mL for the SusCD complex in 0.03% dodecylmaltoside (DDM) detergent. B. SDS-PAGE of the sample peak from panel A shows two prominent bands matching the predicted sizes for SusC and SusD. C. Representative view of grid inside holes with unliganded SusCD complex from which cryo-EM data were collected. D. Representative 2D classes from the top 20 most populated 2D classes of unliganded SusCD complex particles(1).

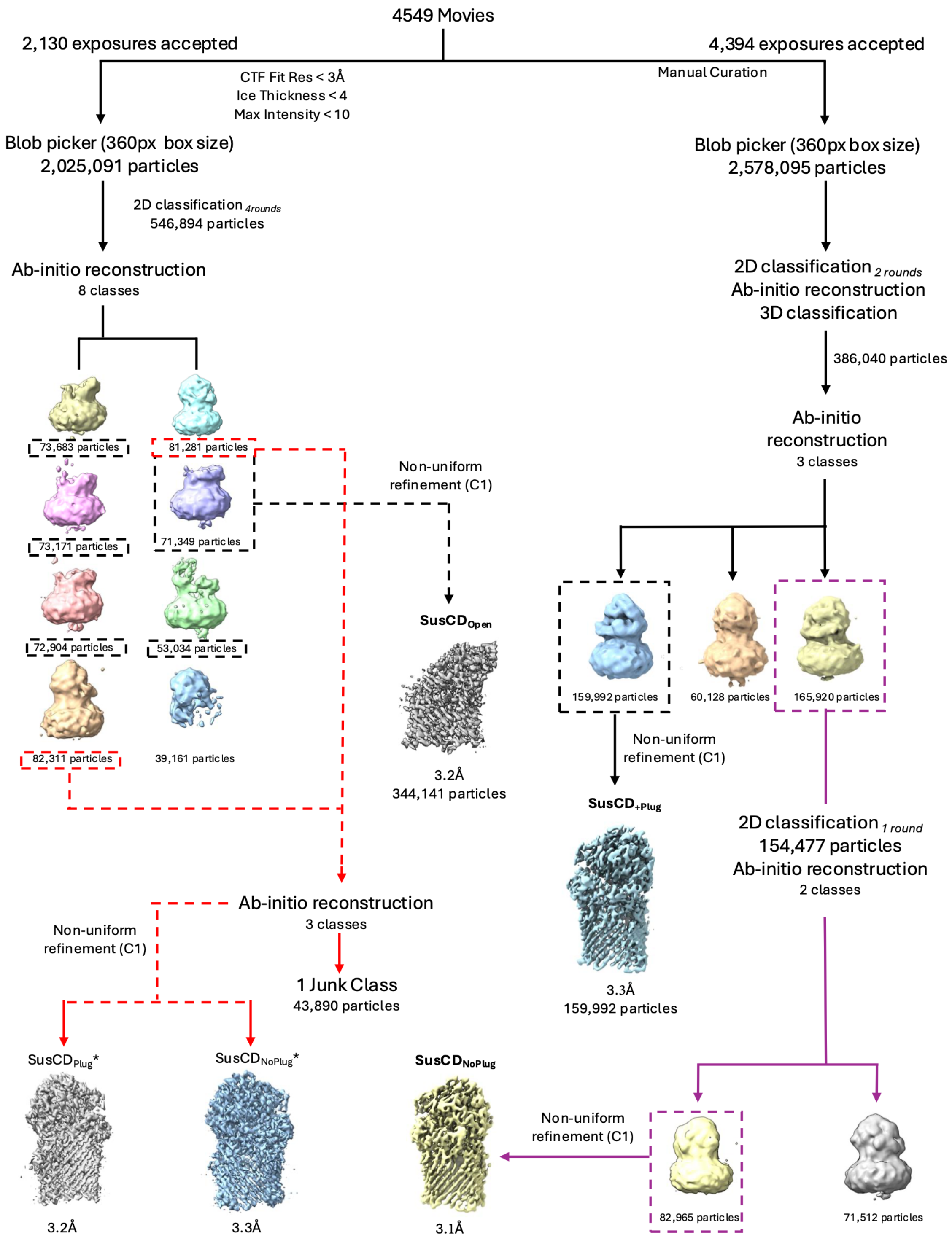


Supplemental Figure 2

**Supplemental Figure 2. Single particle cryo-EM workflow for processing of micrographs into electron density maps for unliganded SusCD**. Starting from a single set of 4,549 movies, two different processing workflows with either strict automatic micrograph curation (left) or manual curation (right) led to the three unliganded SusCD electron density maps. Ab-initio reconstructions are shown with particle selection and curation summary. Maps labeled in bold are final selected maps that were used for model refinement(1).

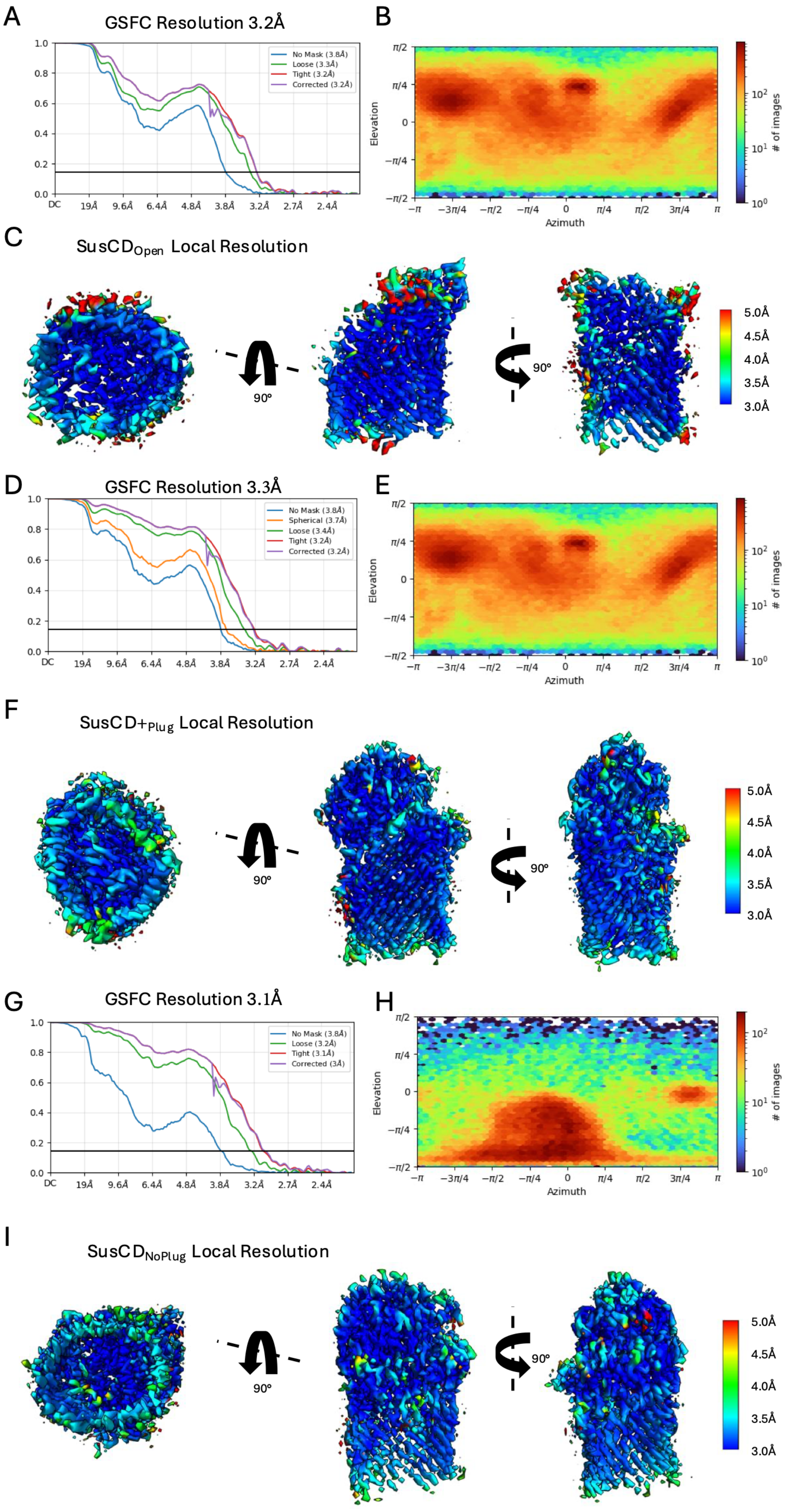


Supplemental Figure 3

**Supplemental Figure 3**. **Gold-standard Fourier shell correlation and local resolution maps for unliganded SusCD complexes.** A. Gold standard Fourier shell correlation (GS-FSC) resolution at a 0.143 cutoff for the final map, showing a global resolution of 3.2Å for the $SusCD_{Open}$ structure. B. Per-particle viewing direction distribution heat map for $SusCD_{Open}$ structure. C. Local resolution map of final refined volume of the $SusCD_{Open}$ reconstruction. Multiple representative angles are shown. D. GS-FSC resolution at a 0.143 cutoff for the final map, showing a global resolution of 3.3Å for the $SusCD_{+Plug}$ structure. E. Per-particle viewing direction distribution heat map for $SusCD_{+Plug}$ structure. F. Local resolution map of final refined volume of the $SusCD_{+Plug}$ reconstruction. Multiple representative angles are shown. G. GS-FSC resolution at a 0.143 cutoff for the final map, showing a global resolution of 3.1Å for the $SusCD_{NoPlug}$ complex. H. Per-particle viewing direction distribution heat map for $SusCD_{NoPlug}$ structure. I. Local resolution map of final refined volume of the $SusCD_{NoPlug}$ reconstruction. Multiple representative angles are shown(1, 2).

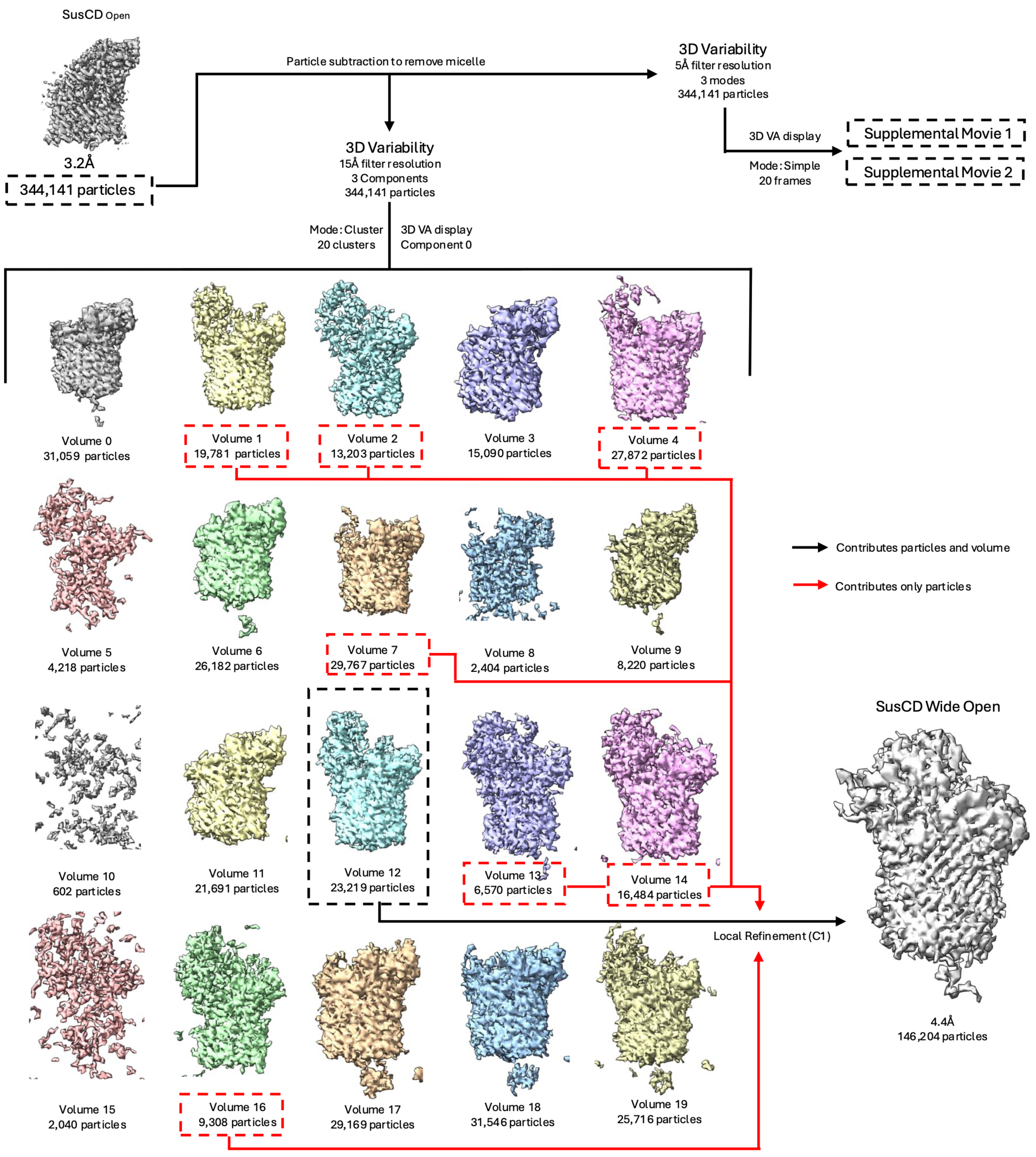


Supplemental Figure 4

**Supplemental Figure 4. 3D variability analysis workflow to capture SusD in an open position**. A workflow for processing particles from the $SusCD_{Open}$ conformation into the refined SusCD open map. Starting from particles of the $SusCD_{Open}$ structure, particle subtraction to remove the micelle was performed and then cryoSPARC 3D variability analysis was performed twice using different filter resolutions. 5Å filter resolution results were visualized as **Movie S1 and S2**. 15Å filter resolution results were clustered into discrete stacks of only component 0 and processed into 3D volumes. The best volume was chosen (indicated by black dashed box) and used as a template for local refinement in combination with secondary particle stacks of similar classes (indicated in red dashed box) to reach a final resolution of 4.4Å with a combined particle stack of 146,204(1–3).

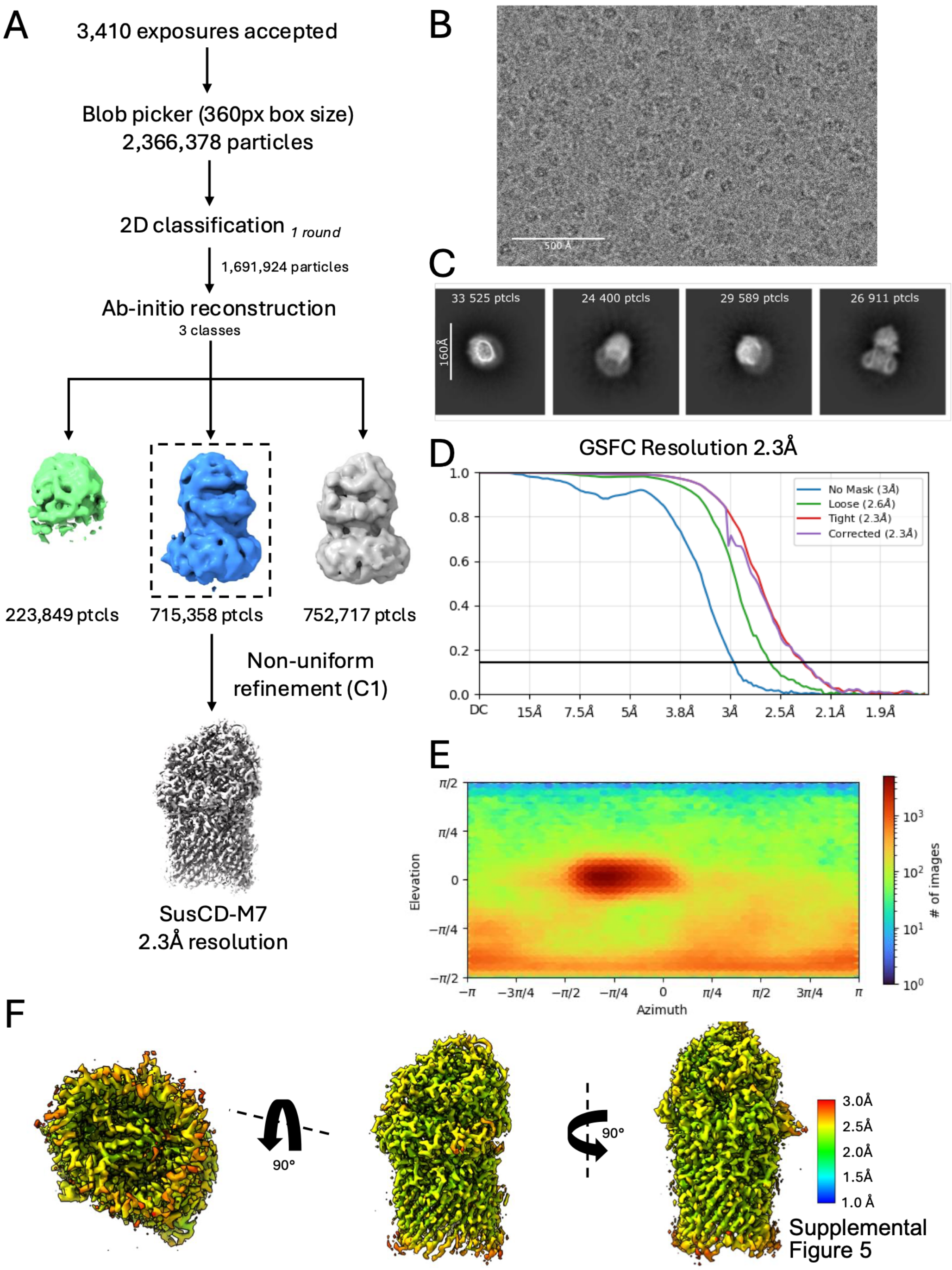


**Supplemental Figure 5. Single particle cryo-EM workflow for processing of micrographs into electron density maps for SusCD with maltoheptaose (SusCD$_{+M7}$)** A. Workflow for processing of micrographs into an electron density map for SusCD-M7. B. Representative view of grid inside holes with SusCD$_{+M7}$ complex from which cryo-EM data was collected. C. Representative 2D classes from the top 20 most populated 2D classes of the SusCD$_{+M7}$ complex particles. D. Gold-standard Fourier shell correlation (GS-FSC) resolution at a 0.143 cutoff for the final map showing a global resolution of 2.3Å for the SusCD$_{+M7}$ structure. E. Per-particle viewing direction distribution heat map for the SusCD$_{+M7}$ structure. F. Local resolution map of final refined volume of the SusCD$_{+M7}$ reconstruction. Multiple representative angles are shown(1, 2).

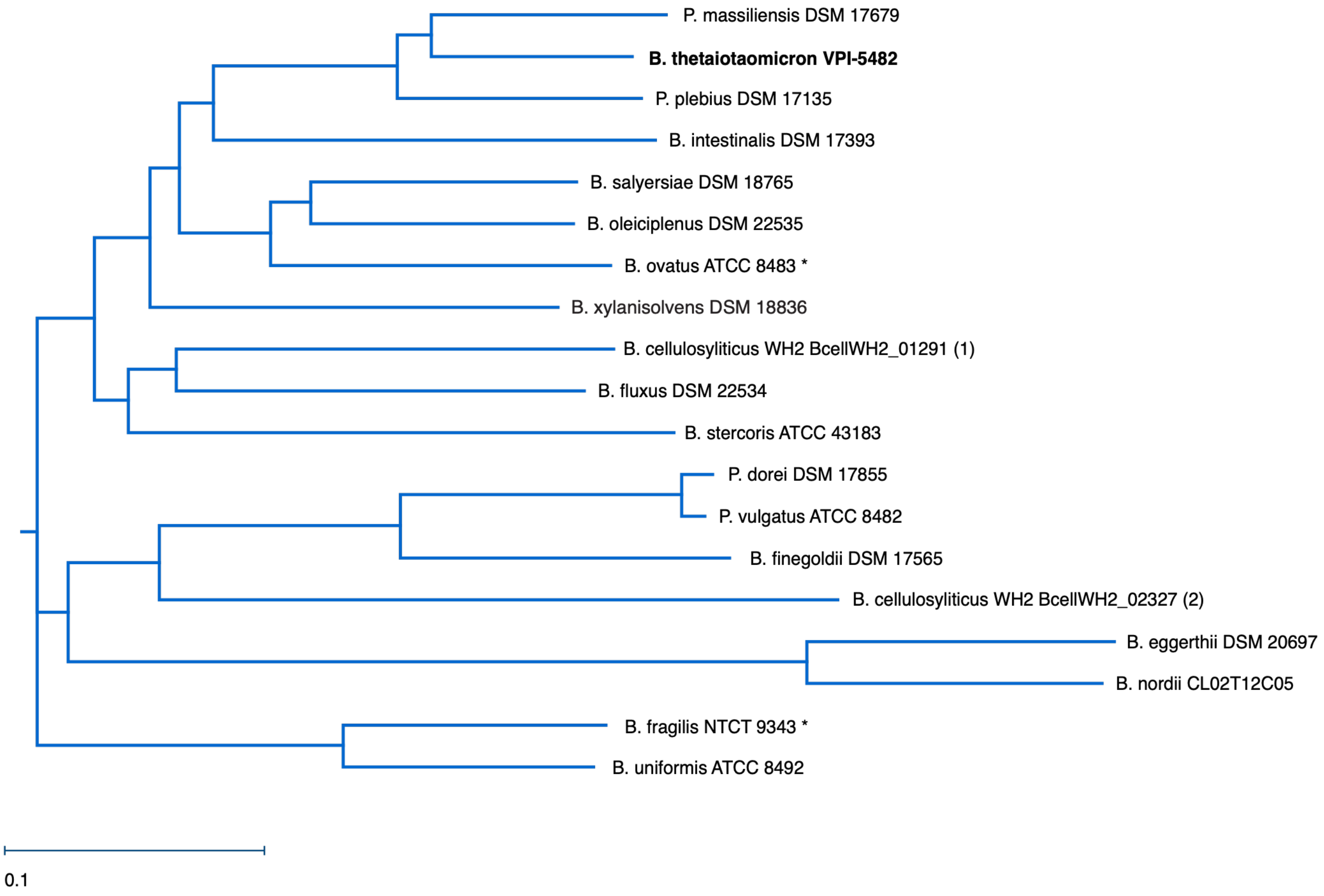


Supplemental Figure 6

**Supplemental Figure 6: Neighbor joining tree of SusC homologs**. Neighbor joining tree obtained from the multisequence alignment show in Supplemental Figure 7. Sequences were taken from Sus PUL in *Bacteroides* type strains that grow on starch(4). Bt SusC is in bold, and * indicates the two sequences for which AlphaFold3 predictions of monomer and dimer assemblies were created as described in the text. Tree created in MegAlignPro within DNAstar (5). Protein sequences: *Phocaeicola massiliensis* HMPREF1534_02829, *Bacteroides thetaiotaomicron* BT_3702, *Phocaeicola plebius* BACPLE_02370, *Bacteroides intestinalis* BACINT_03131, *Bacteroides salyersiae* HMPREF1532_02495, *Bacteroides oleiciplenus* WP_009132665, *Bacteroides ovatus* Bovatus_03807, *Bacteroides xylanisolvens* BXY_47640, *Bacteroides cellulosyliticus* BcellWH2_01291, *Bacteroides fluxus* HMPREF9446_00914, *Bacteroides stercoris* BACSTE_01657, *Phocaeicola dorei* WP_008674305, *Phocaeicola vulgatus* BVU_1380, *Bacteroides finegoldii* BACFIN_07563, *Bacteroides cellulosyliticus* BcellWH2_02327, *Bacteroides eggerthii* BACEGG_01900, *Bacteroides nordii* HMPREF1068_03059, *Bacteroides fragilis* BF3146, *Bacteroides uniformis* BACUNI_01209.

**Supplement Fig 7: Multisequence alignment of SusC Homologs**. Alignment was created via CLUSTAL W and visualized in MegAlignPro (5, 6). Structural features are indicated above the BtSusC ruler, M7 binding site one residues are indicated with red arrows and site two with blue arrows. The intracellular loop contributing to synthetic SusC dimer clashing labeled as “dimer clash”. Protein sequences: *Phocaeicola massiliensis* HMPREF1534_02829, *Bacteroides thetaiotaomicron* BT_3702, *Phocaeicola plebius* BACPLE_02370, *Bacteroides intestinalis* BACINT_03131, *Bacteroides salyersiae* HMPREF1532_02495, *Bacteroides oleiciplenus* WP_009132665, *Bacteroides ovatus* Bovatus_03807, *Bacteroides xylanisolvens* BXY_47640, *Bacteroides cellulosyliticus* BcellWH2_01291 (sequence labeled 1), *Bacteroides fluxus* HMPREF9446_00914, *Bacteroides stercoris* BACSTE_01657, *Phocaeicola dorei* WP_008674305, *Phocaeicola vulgatus* BVU_1380, *Bacteroides finegoldii* BACFIN_07563, *Bacteroides cellulosyliticus* BcellWH2_02327 (sequenced labeled 2), *Bacteroides eggerthii* BACEGG_01900, *Bacteroides nordii* HMPREF1068_03059, *Bacteroides fragilis* BF3146, *Bacteroides uniformis* BACUNI_01209.

Supplemental Figure 7

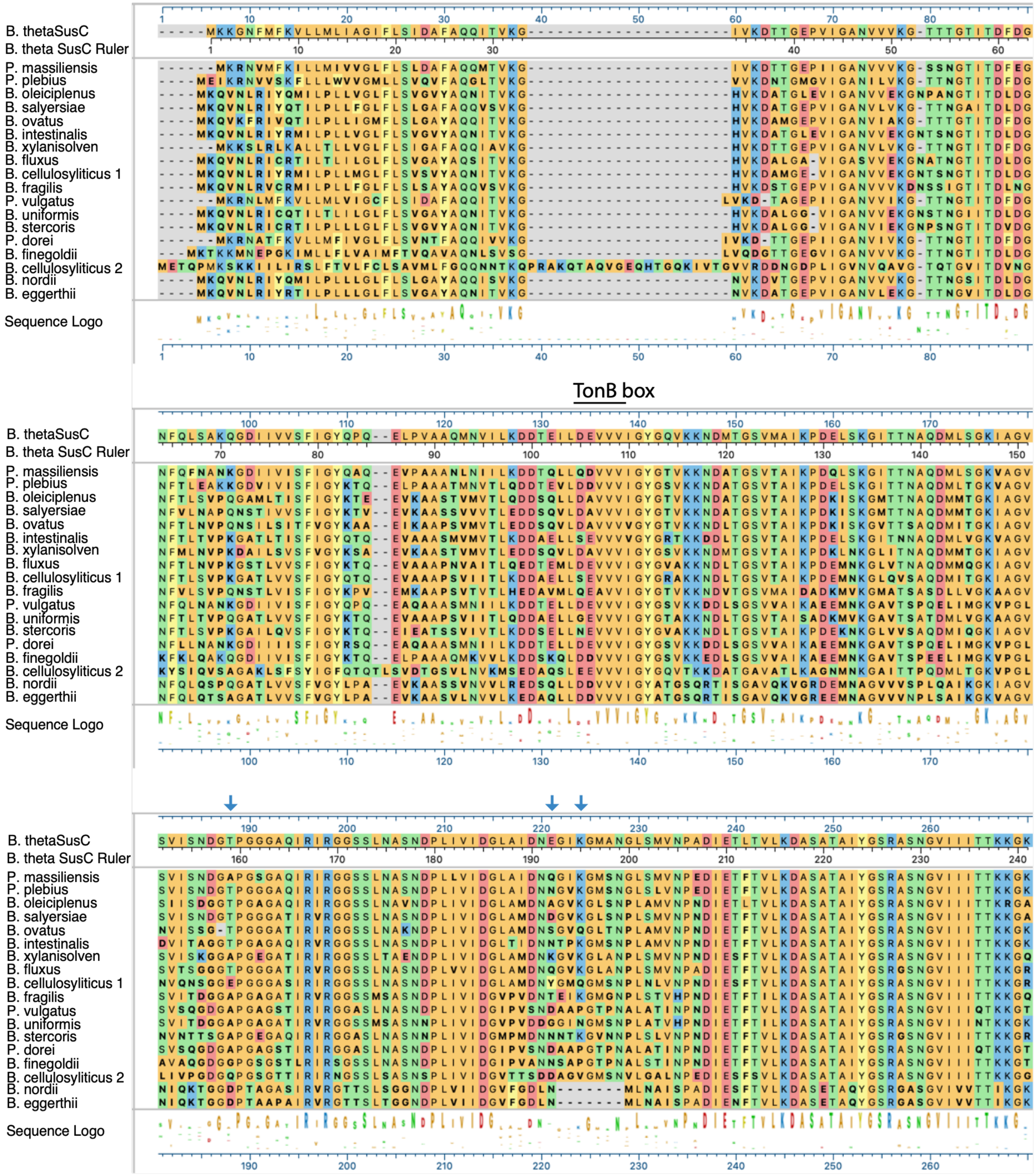

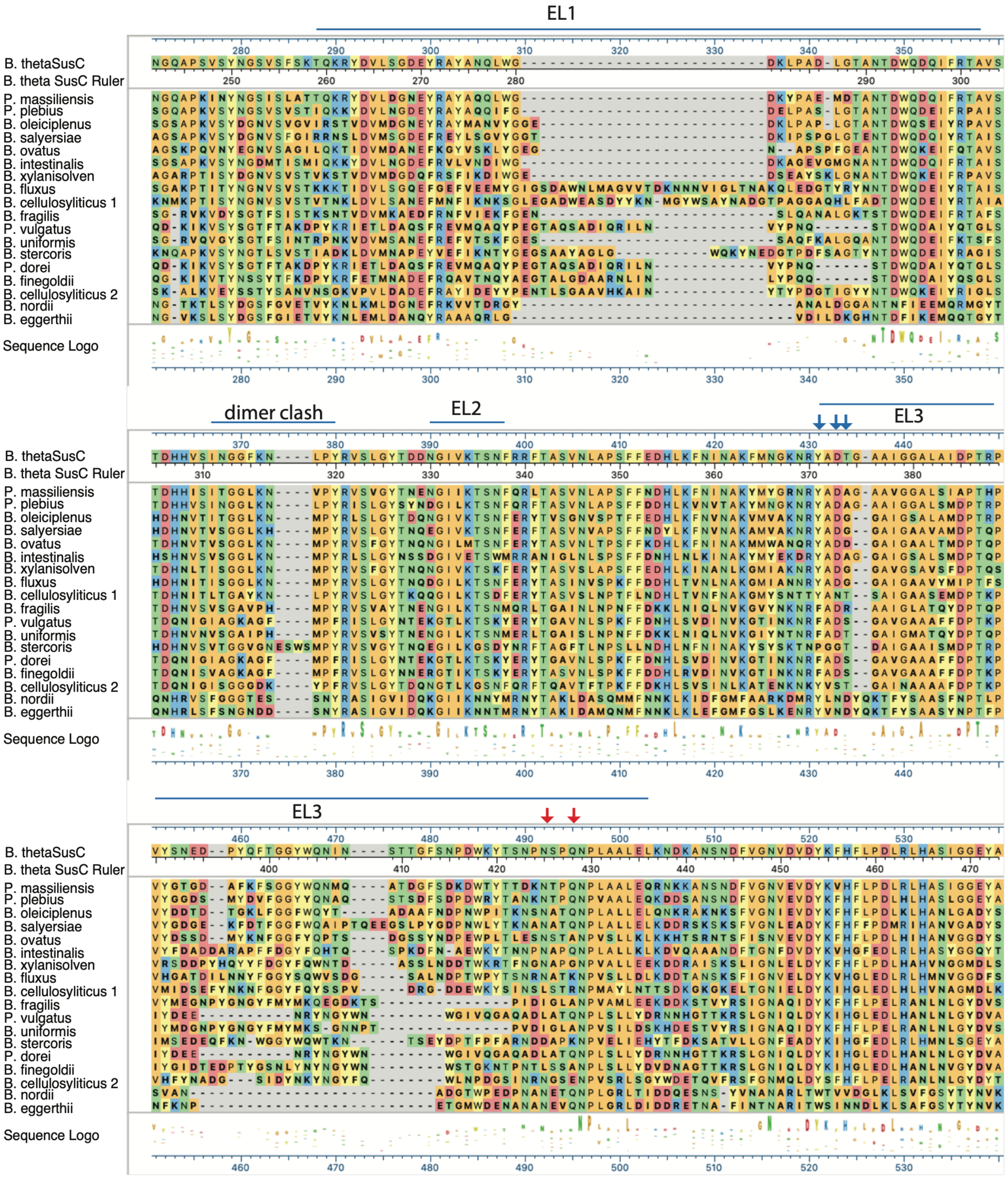
EL1
dimer clash
EL2
EL3
B. thetaSusC
B. theta SusC Ruler
P. massiliensis
P. plebius
B. oleiciplenus
B. salyersiae
B. ovatus
B. intestinalis
B. xylanisolven
B. fluxus
B. cellulosyliticus 1
B. fragilis
P. vulgatus
B. uniformis
B. stercoris
P. dorei
B. finegoldii
B. cellulosyliticus 2
B. nordii
B. eggerthii
Sequence Logo

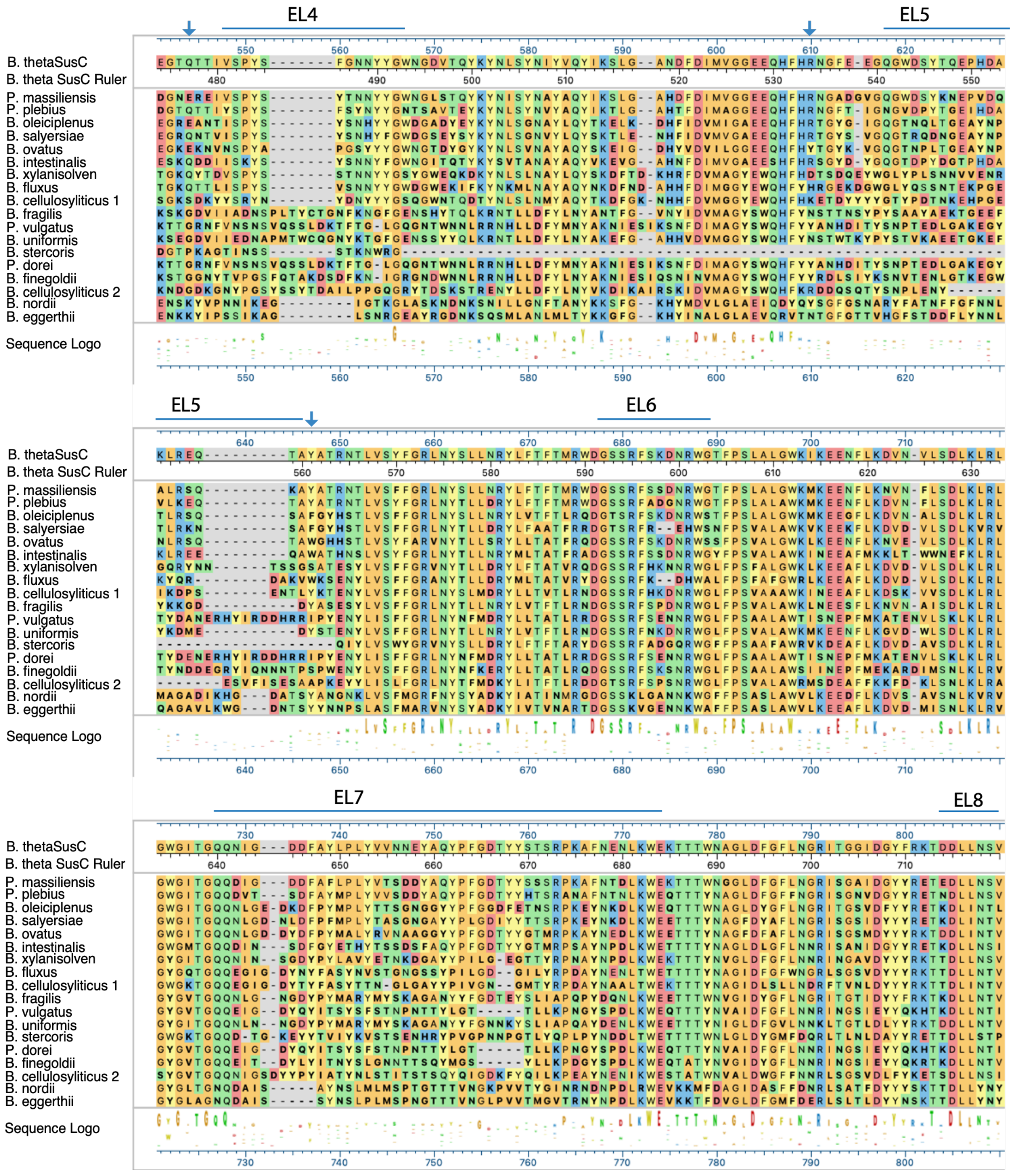
EL4
EL5
EL6
EL7
EL8
B. thetaSusC
B. theta SusC Ruler
P. massiliensis
P. plebius
B. oleiciplenus
B. salyersiae
B. ovatus
B. intestinalis
B. xylanisolven
B. fluxus
B. cellulosyliticus 1
B. fragilis
P. vulgatus
B. uniformis
B. stercoris
P. dorei
B. finegoldii
B. cellulosyliticus 2
B. nordii
B. eggerthii
Sequence Logo

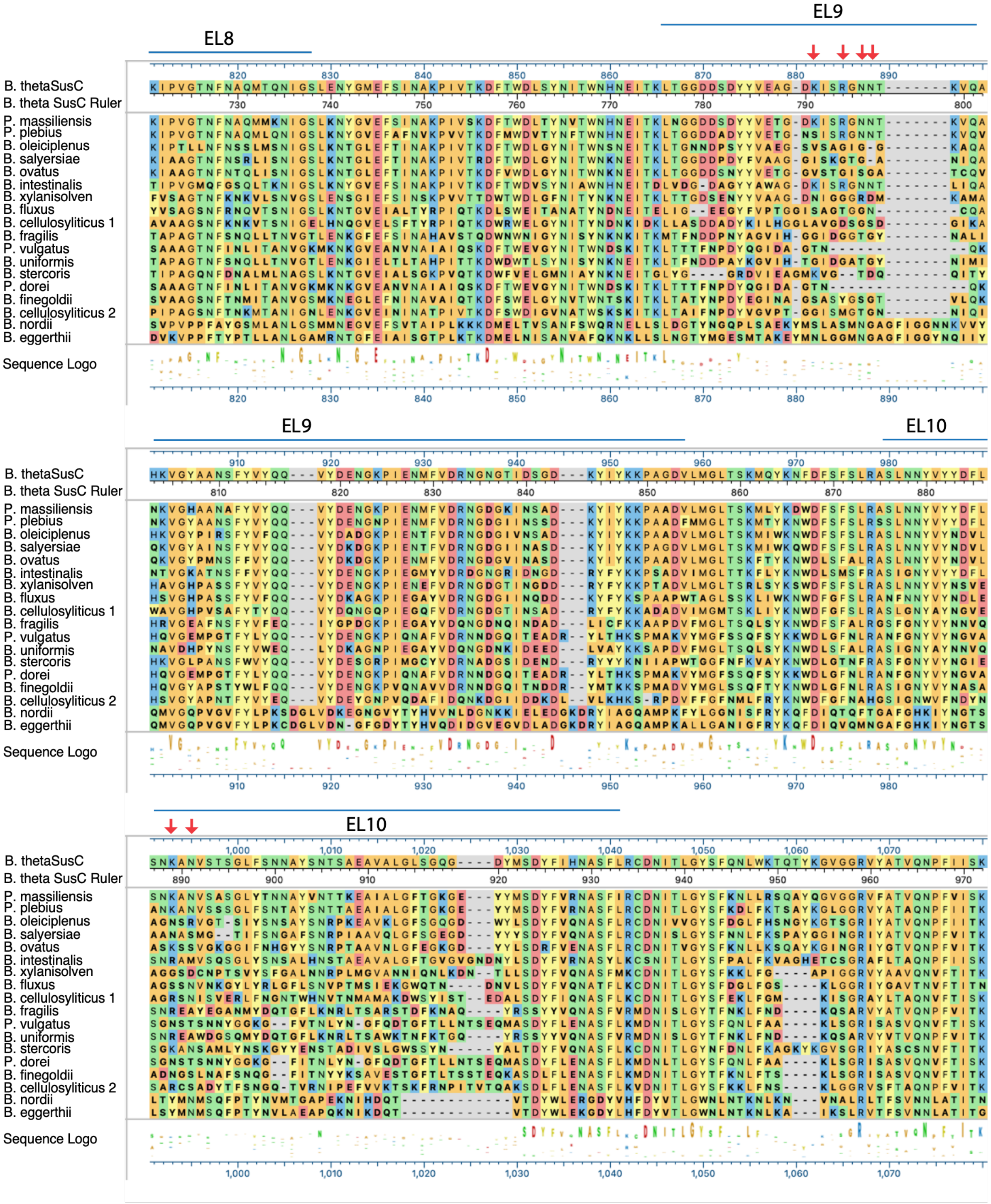
EL8
EL9
EL10
B. thetaSusC
B. theta SusC Ruler
P. massiliensis
P. plebius
B. oleiciplenus
B. salyersiae
B. ovatus
B. intestinalis
B. xylanisolven
B. fluxus
B. cellulosyliticus 1
B. fragilis
P. vulgatus
B. uniformis
B. stercoris
P. dorei
B. finegoldii
B. cellulosyliticus 2
B. nordii
B. eggerthii
Sequence Logo

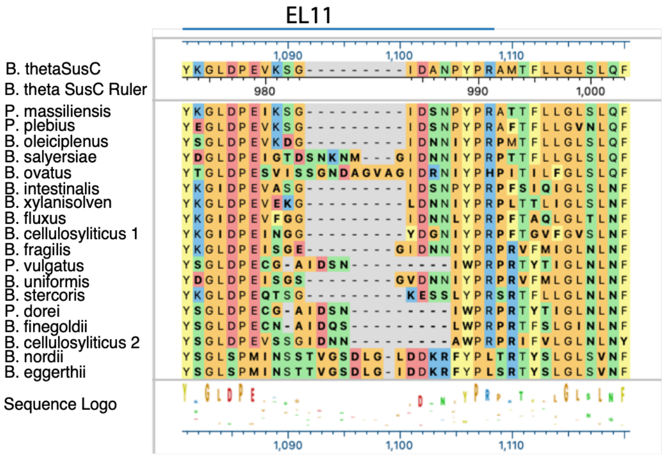
EL11
1,090
1,100
1,110
B. thetaSusC
B. theta SusC Ruler
980
990
1,000
P. massiliensis
P. plebius
B. oleiciplenus
B. salyersiae
B. ovatus
B. intestinalis
B. xylanisolven
B. fluxus
B. cellulosyliticus 1
B. fragilis
P. vulgatus
B. uniformis
B. stercoris
P. dorei
B. finegoldii
B. cellulosyliticus 2
B. nordii
B. eggerthii
Sequence Logo
1,090
1,100
1,110

| Table S1: Microscopy and Model Statistics | | | | | |
|---|---|---|---|---|---|
| | $SusCD_{+M7}$ | $SusCD_{Open}$ | $SusCD_{+Plug}$ | $SusCD_{NoPlug}$ | $SusCD_{3DVA}$ |
| **Data collection** | | | | | |
| Instrument | Titan Krios G3 | Titan Krios G3 | Titan Krios G3 | Titan Krios G3 | Titan Krios G3 |
| Detector | Gatan K3 | Gatan K3 | Gatan K3 | Gatan K3 | Gatan K3 |
| Magnification | 105,000x | 81000X | 81000X | 81000X | 81000X |
| Voltage (kV) | 300 | 300 | 300 | 300 | 300 |
| Total electron dose ($e^-/Å^2$) | 55 | 60 | 60 | 60 | 60 |
| Defocus range (μm) | -0.4 to -3.5 | -0.7 to -3.4 | -1.2 to -3.8 | -1.2 to -3.8 | -0.7 to -3.4 |
| Pixel size (Å) | 0.83956 | 1.081* | 1.0672 | 1.0672 | 1.0672 |
| Processing software | CryoSPARC | CryoSPARC | CryoSPARC | CryoSPARC | CryoSPARC |
| Initial particles | 2,366,378 | 2,025,091 | 2,578,095 | 2,578,095 | 2,025,091 |
| Final particles | 715,358 | 344,141 | 159,992 | 82,965 | 80,677 |
| Map sharpening β-factor | 81.4 | 108.1 | 90.3 | 65.7 | 418.6 |
| Map resolution (Å) | 2.3 | 3.2 | 3.3 | 3.1 | 4.4 |
| FSC threshold | 0.143 | 0.143 | 0.143 | 0.143 | 0.143 |
| **Validation** | | | | | |
| MolProbity score | 1.45 | 2.35 | 2.53 | 2.43 | - |
| MolProbity clashscore | 3.77 | 6.46 | 11.76 | 12.38 | - |
| Rotamer outliers (%) | 1.68 | 5.43 | 4.81 | 4.06 | - |
| $C_\beta$ deviations (%) | NA | NA | NA | NA | - |
| **R.M.S. deviations** | | | | | |
| Bond lengths (Å) | 0.011 | 0.004 | 0.005 | 0.004 | - |
| Bond angles (°) | 0.804 | 0.686 | 1.045 | 0.663 | - |
| **Ramachandran Plot** | | | | | |
| Favored (%) | 97.45 | 93.24 | 93.48 | 94.82 | - |
| Allowed (%) | 2.55 | 6.64 | 6.37 | 5.18 | - |
| Outliers (%) | 0 | 0.12 | 0.15 | 0 | - |

* Phenix.magref was used to refine pixel spacing against the AlphaFold3 predicted structure of SusC prior to model fitting and refinement.

| **Table S2: Primers for *susD-6xhis* allele in pExchange** | | |
|---|---|---|
| Primer name | Sequence (5' – 3') | Use |
| **SusCup-Sal1** | GAAGATAACATTCGA**gtcgac** CACCTGGGACCTCAGCTATAAC | *susC* forward primer, anneals ~750 bp up from gene end |
| **SusDhis-rev** | ATGATGATGGTGGTGATGAGCTGCTGC TTTATAGCCTTCATTTTGTGAC | *susD* reverse primer that includes 6x His tag and 3x Ala linker for SusD |
| **SusDhis-for** | GCAGCAGCTCATCACCACCATCATCAT TAACCAAGAGTTCATCCTTATATAAAAG | *susD* forward primer anneals to new 3x Ala/6x His fusion and intergenic region between *susD* and *susE* |
| **SusEdown-Xba1** | CGCGGTGGCGGCCGC**tctaga** CTGACTCCTGCTCCGGCTTTG | *susE* reverse primer, anneals to middle of SusE gene |

Region that anneals to pExchange is in red and restriction site is in lower case bold.

**Movie S1:** 3D variability movie generated from CryoSPARC using SusCD open conformation particles. The reaction coordinate demonstrates changes in SusD from an open to more closed position as appearance of density for the NTE is also observed(3).

**Movie S2:** 3D variability movie generated from CryoSPARC using SusCD open conformation particles. The reaction coordinate demonstrates changes in SusD from a closed to more open position as changes in the shape of the NTE are also observed(3).

**Movie S3:** Morph between $SusCD_{Open}$ and $SusCD_{+Plug}$ maps using ChimeraX demonstrating the changes in the open and closed conformations of SusCD such as when binding ligand(2).

1. Punjani, A., Rubinstein, J. L., Fleet, D. J., and Brubaker, M. A. (2017) CryoSPARC: Algorithms for rapid unsupervised cryo-EM structure determination. *Nat. Methods*. **14**, 290–296

2. Meng, E. C., Goddard, T. D., Pettersen, E. F., Couch, G. S., Pearson, Z. J., Morris, J. H., and Ferrin, T. E. (2023) UCSF ChimeraX: Tools for structure building and analysis. *Protein Science*. 10.1002/pro.4792

3. Punjani, A., and Fleet, D. J. (2021) 3D variability analysis: Resolving continuous flexibility and discrete heterogeneity from single particle cryo-EM. *J. Struct. Biol.* 10.1016/j.jsb.2021.107702

4. Pudlo, N. A., Urs, K., Crawford, R., Pirani, A., Atherly, T., Jimenez, R., Terrapon, N., Henrissat, B., Peterson, D., Ziemer, C., Snitkin, E., and Martens, E. C. (2022) Phenotypic and Genomic Diversification in Complex Carbohydrate-Degrading Human Gut Bacteria. *mSystems*

5. Burland, T. G. *DNASTAR's Lasergene Sequence Analysis Software*

6. Thompson, J. D., Gibson, Toby. J., and Higgins, D. G. (2003) Multiple Sequence Alignment Using ClustalW and ClustalX. *Curr. Protoc. Bioinformatics*. 10.1002/0471250953.bi0203s00
