## Supplementary material for "Structural analyses of a heterodimeric SusCD complex captures intermediate states of maltooligosaccharide transport": Author Contributions

**Michael C. Cadigan**: Conceptualization, Methodology and Formal Analysis, Writing- Original Draft, Writing – Review and Editing, Data Curation, Data Visualization; **Nicole Rivera Fuentes**: Methodology and Formal Analysis, Data Visualization, **Wilhelm Salmen**: Methodology and Formal Analysis, Data Visualization, Writing – Review and Editing; **Jacquelyn R. Roberts**: Methodology and Formal Analysis; **Brandon Ruotolo**: Methodology and Formal Analysis, Data Visualization, Writing – Review and Editing, Supervision, Funding Acquisition; **Melanie D. Ohi**: : Conceptualization, Methodology and Formal Analysis, Writing- Original Draft, Writing – Review and Editing, Data Curation, Data Visualization, Supervision, Funding Acquisition. **Nicole M. Koropatkin:** Conceptualization, Methodology and Formal Analysis, Writing- Original Draft, Writing – Review and Editing, Data Curation, Data Visualization, Supervision, Funding Acquisition.
